# Blood Pressure Regulates Functional Coupling of L-Type Ca^2+^ Channels: Reimaging the Foundation of Cerebral Blood Flow Control

**DOI:** 10.64898/2026.03.02.708544

**Authors:** Galina Yu. Mironova, Miguel Martín-Aragón Baudel, Chryso Lambride, Sanjay R. Kharche, David A. Steven, Jonathan Lau, Keith W. MacDougall, Melfort Boulton, Franca Schmid, Manuel F. Navedo, Donald G. Welsh

**Author notes:** **Correspondence:** Galina Yu. Mironova, Ph.D., Robarts Research Institute, Dept Physiology & Pharmacology, 1151 Richmond St, University of Western Ontario, London, ON N6A 3K7, or Donald G. Welsh, Ph.D., Robarts Research Institute, Dept Physiology & Pharmacology, 1151 Richmond St, University of Western Ontario, London, ON N6A 3K7. **Author Contributions: GYM** and **DGW** designed the study; **GYM –** performed experiments, conducted data analysis and prepared the figures (except super-resolution experiments and **Fig. 9**), wrote the original and revised manuscript with inputs from **SRK**, **MFN & DGW**; **SRK** – developed an in-house algorithm (Markov chains) to analyze single-channel activity; **MMB&MFN** – performed super resolution microscopy experiments and analysis; **MFN** –provided access to S1928A mutant mouse; **DAS, JL**, **KWM&MB** – clinicians essential for procurement of human tissue; **CL & FS** – developed computer blood flow model and conducted in silico simulations; **MFN**,**DGW** – conducted general edits to the final version of the manuscript. All authors approved the final version of the manuscript. **Competing Interest Statement** Authors claim no competing interests.

## Abstract

The myogenic response is the key autoregulatory mechanism setting cerebral blood flow and its mechanistic foundation is intimately tied to depolarization and the voltage gating of L-type Ca^2+^ channels (Ca_V_1.2). While critical, this study argues for an additional mechanism, that of pressure itself enhancing Ca_V_1.2 activity via cooperative gating and perimembrane trafficking of channel’s subunits. These novel insights were pursued at the cell level using patch-clamp electrophysiology and advanced microscopy, and then functionally in pressurized arteries through measures of tone and intracellular [Ca^2+^]_i_. Key findings were confirmed in mutant mice with disrupted functional coupling and translated into arteries procured from human brain tissue. From cerebral blood flow simulations of semi-realistic microvascular networks, we predict that loss of this alternative mechanism leads to maldistribution of brain blood flow and potentially a diminishment of cognitive function. This study reveals previously unrecognized pressure-sensitive Ca_V_1.2 regulatory mechanism that advances understanding of cerebral blood flow.

**Significance:** Blood pressure sets base arterial constriction - a response critical for blood flow control in brain. This response is tied to Ca_V_1.2 channels and their presumptive and exclusive activation by voltage, reasoning now under great scrutiny. We establish herein with advance methods, a second mode of Ca_V_1.2 pressure regulation, that of enhanced functional cooperativity among neighboring channels. This novel mechano-response is tied to PKCα and its ability to set channel phosphorylation and Ca_V_1.2 trafficking. Ca_V_1.2 pressure regulation was observed in human tissues and its disruption (mutant mice) impaired myogenic tone in the presence of preserved voltage control. Cerebral microvascular modeling highlights that losing this mechanism destabilizes blood flow distribution in brain, the knock-on effect being comprised cognitive function.

## Introduction

William Bayliss first described arterial constriction to elevated intravascular pressure, the so called “myogenic response” now viewed as foundational to the control of blood pressure, base blood flow and capillary pressure (1). What is unique with this response is its near complete dependence on L-type Ca^2+^ channels to elevate cytosolic [Ca^2+^] in the service of myosin light chain phosphorylation. It’s typically assumed that myogenic depolarization is the sole factor responsible for opening a resident pool of L-type Ca^2+^ channels (2). While a bedrock principle, we query herein if pressure modulates L-type Ca^2+^ channels through other means so to ensure greater Ca^2+^ influx per mV depolarization, perhaps in countenance to elevated wall stress.

L-type Ca^2+^ channels are comprised of an α1 pore-forming subunit (Ca_V_1.2), along with auxiliary proteins (α2, β, *δ*, and γ) that control channel gating/kinetics and trafficking (3, 4). In vascular smooth muscle, one regulatory aspect that has received concerted attention is functional coupling - a biophysical state whereby two or more Ca_V_1.2 subunits interact via their C-termini augmenting their activity (4–6). Literature reveals that functional coupling is dynamically regulated by protein kinases inducing 1) Ca_V_1.2 C-terminus phosphorylation (6), and potentially 2) perimembrane trafficking (7). When disrupted by disease, for example diabetes and hypertension, arterial dysfunction ensues as does blood flow maldistribution (8–10). But what of homeostatic physiology: could functional coupling be intimately tied to myogenic tone development and if so, is it under protein kinase control?

This study examined whether and by what mechanism intravascular pressure facilitates the functional coupling of L-type Ca^2+^ channels in service of myogenic tone development. We interrogated rodent cerebral vasculature using patch-clamp electrophysiology, super-resolution microscopy and pressure myography. We report herein that mechano-stimuli do indeed augment functional coupling, through PKCα regulation and that this overall phenomenon, tied to dynamic channel clustering, is essential to pressure-induced tone development. Key observations were replicated in human cerebral arteries and note that these responses were compromised in transgenic mice with disrupted Ca_V_1.2-S1928 phosphorylation across the physiological pressure range. Ensuing hemodynamic modeling reveals that disruption of functional coupling leads to marked blood flow maldistribution, with particular brain regions being over and under perfused. We conclude pressure-induced regulation of L-type Ca^2+^ channels extend beyond voltage control and is synergistically dependent on channel clustering and gating cooperativity that ensues.

## Results

### Modulation of L-Type Ca^2+^ Channel Activity by Mechano-stimuli

We began our examination by monitoring whole-cell Ba^2+^ current in isoosmotic (∼300 mOsm) and hypoosmotic (∼230 mOsm) bath solutions, the latter is a mechano-stimulus that swells/stretches smooth muscle cells imitating the increase of blood pressure (**Fig. 1A, top**). We chose rat cerebral arterial myocytes to match the current work with experiments sporadically found in the historical literature (11–15). In alignment, our swelling perturbation increased the Ca_V_1.2 (∼30% at 10mV) current without impacting voltage dependency of activation and steady-state inactivation (**Fig. 1B & 1C**). In contrast, the whole-cell Ca_V_3.1 current, a secondary source of contractile Ca^2+^, failed to respond to the swelling protocol (**Suppl. Fig. S1**). Swelling’s impact on Ca_V_1.2 could in theory originate from enhanced functional coupling, a process mediated in part by PKC or PKA (**Fig. 1A, bottom**). Consistent with theory, PKC inhibition (Calphostin C) abolished the swelling-induced Ca_V_1.2 current (**Fig. 1D**), whereas blockade of AKAP-PKA interaction (st-Ht31) had no significant effect (**Fig. 1E**). Note, both pharmacological interventions augmented baseline Ca_V_1.2 current (**Fig. 1F, grey bars**), the strongest effect mediated by Calphostin C.

**Figure 1.**
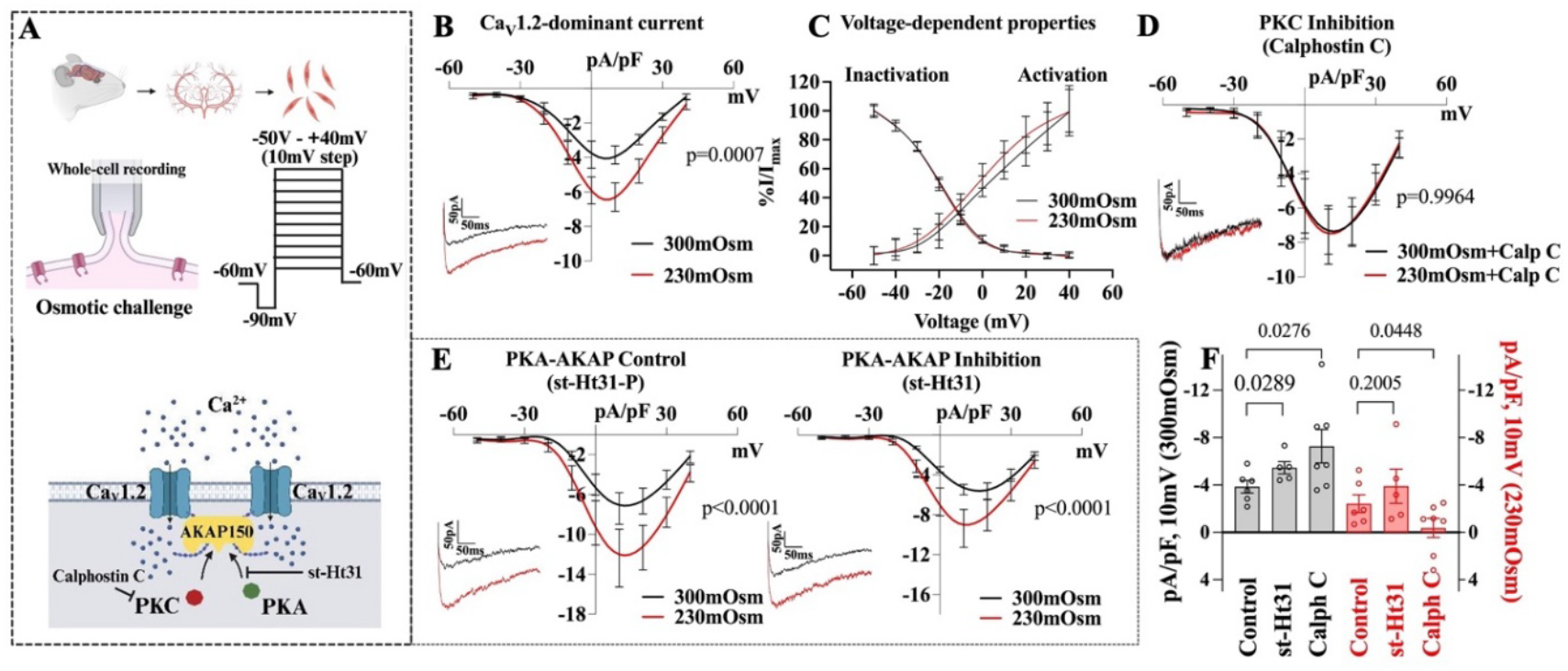
Mechano-stimulation of rat cerebral arterial smooth muscle cells increases whole-cell Ca_V_1.2 current. **A(top)**, Schematic of cerebral arterial smooth muscle cell isolation and whole-cell patch-clamp protocol. To simulate membrane stretch by intravascular pressure, cells were exposed to a hypoosmotic bath solution (300-to-230 mOsm). **A (bottom)**, Schematic of functionally coupled Ca_V_1.2 with AKAP tethering PKA & PKC to the C-terminus. Protein kinase regulation was disrupted by the application of Calphostin C (300 nM, pan PKC inhibitor) and st-Ht31 (10μM, disrupts AKAP-PKA interaction). **B**, Whole-cell Ca_V_1.2 current density is higher following exposure to a hypoosmotic challenge (n=6 cells from 6 animals, two-way ANOVA, Sidak’s multiple comparison test, p shown for the peak value). **C**, Voltage dependency of Ca_V_1.2 activation (n=6 cells from 6 animals) and steady-state inactivation (n=13 cells from 13 animals) is unaffected by hypoosmotic challenge. **D**, PKC inhibition by Calphostin C (300 nM) abolishes the hypoosmotic-induced increase in whole-cell Ca_V_1.2 current (n=7 cells from 7 animals, two-way ANOVA, Sidak’s multiple comparison test, no significant difference, p shown for the peak value). **E**, Disrupting PKA-AKAP interaction with st-Ht31 (10 µM) did not affect the hypoosmotic-induced increase in whole-cell Ca_V_1.2 current; the control peptide st-Ht31-P (10 µM) was equally ineffective (left – control, right – experiment; n=5 cells from 5 animals, two-way ANOVA, Sidak’s multiple comparison test, p shown for the peak values). **F**, Summary of whole-cell data demonstrating that PKC inhibition, but not AKAP-PKA, significantly impacts pressure-sensitive current (red bars, unpaired Welch’s t-test with a one-tailed hypothesis). Note, the PKC inhibitor, more so than the PKA-AKAP affected baseline current at 300 mOsm (grey bars, unpaired Welch’s t-test with a one-tailed hypothesis); n of cells and animals same as in corresponding whole-cells experiments.

We next pursued a definitive single-channel assessment of Ca_V_1.2 coupling properties using smooth muscle cells isolated from mouse cerebral arteries. We specifically employed cell-attached patch-clamp to quantify activity at rest and following negative pressure application via the patch pipette (-20 mmHg, **Fig. 2 flowchart**). Compared to base conditions, single-channel Ca_V_1.2 activity markedly increased with pressurization ast noted in traces (**Fig. 2A**) and quantitative analysis; the number of channels opening (**Suppl. Figure S9)**, open probability (**Fig. 2B**), coupling frequency (**Suppl. Figure S9**), and coupling strength (**Fig. 2C**). Controls confirmed that single channel activity was inhibited by Nifedipine (L-type Ca^2+^ channel blocker, **Suppl. Fig. S2**). Pretreatment of cells with a broad PKC (Calphostin C, **Fig.2D**) or PKCα selective (Go-6976, **Fig. 2G**) inhibitors suppressed the pressure-induced increase in number of channels opening (**Suppl. Figure S8**), open probability (**Fig. 2E&H**), coupling frequency (**Suppl. Figure S8**), and coupling strength (**Fig. 2F&I**). Akin to whole-cell measures, baseline activity increased following PKC inhibition, including the number of channels opening (**Suppl. Figure S8**). The latter aligns with pressure potentially enhancing functional coupling by promoting Ca_V_1.2 subunit trafficking to the plasma membrane.

**Figure 2.**
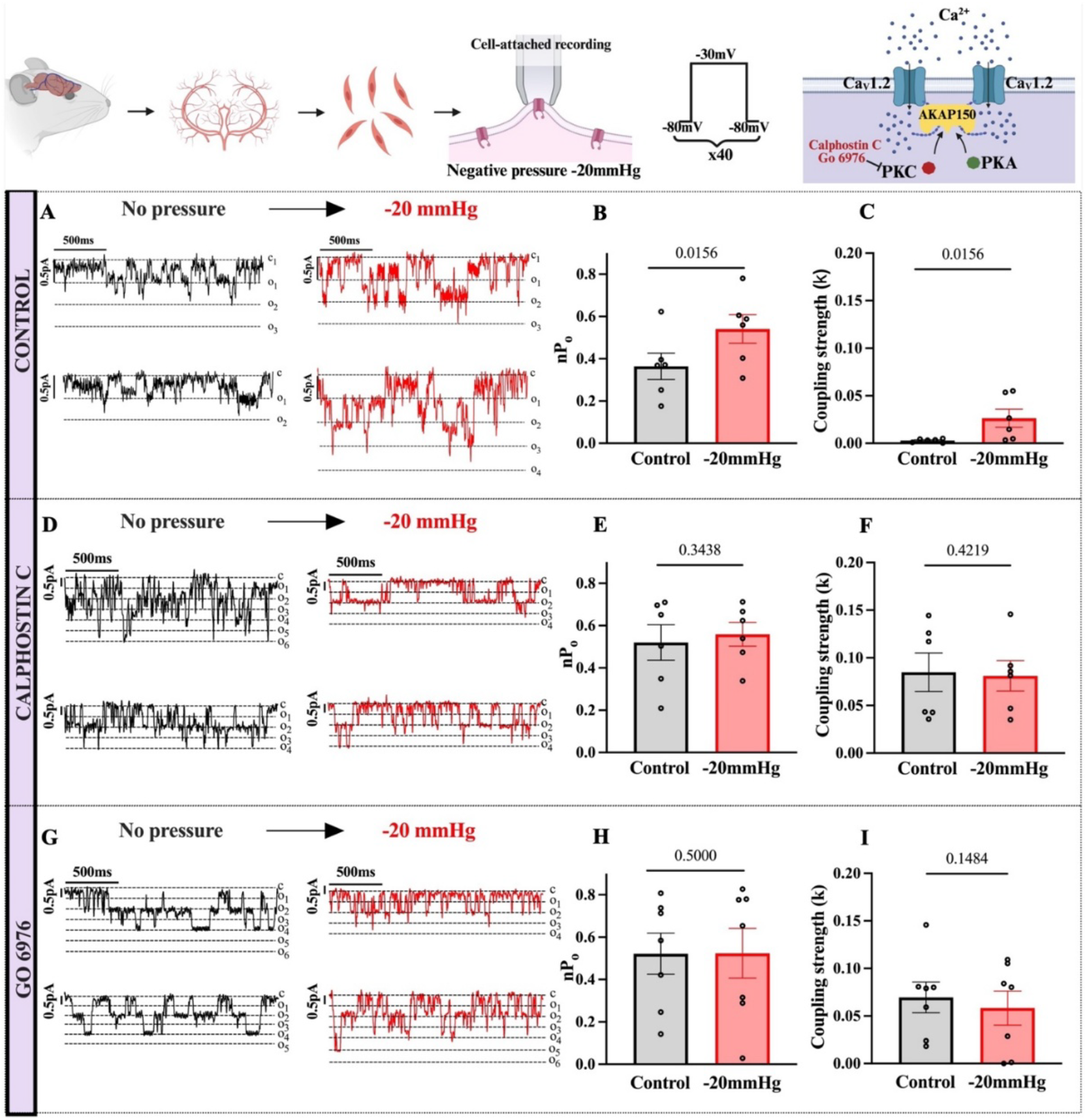
Mechano-stimulation increases Ca_V_1.2 single-channel activity in mouse cerebral arterial smooth muscle cells and depends on PKC. Top chart shows schematic of cerebral arterial smooth muscle cells isolation and the ensuing cell-attached single-channel recording protocol. To simulate membrane stretch, negative pressure (-20 mmHg) was applied via a patch pipette. Representative traces of cell-attached, single-channel recordings of Ca_V_1.2 activity prior to and following negative pipet pressure application (-20 mmHg) under control conditions (**A**), and in presence of general PKC inhibitor Calphostin C (300 nM) (**D**), and specific PKC inhibitor Go 6976 (100 nM) (**G**). The statistical analysis of single-channel data demonstrated that negative pressure application increased an open probability and coupling strength of Ca_V_1.2 (**B-C** respectively, n=6 cells from 6 animals, Wilcoxon matched-pairs signed-rank test with a one-tailed hypothesis). Application of PKC inhibitors Calphostin C or Go 6976 both inhibited open probability and coupling strength increase of Ca_V_1.2 (**E-F** and **H-I** respectively, n=6 cells from 6 animals for Calphostin C/n=7 cells from 7 animals for Go 6976, Wilcoxon matched-pairs signed-rank test with a one-tailed hypothesis).

### Mechano-Stimuli and Ca_V_1.2 Trafficking

Ca_V_1.2 subunits reside in caveolae and closely associated perimembrane vesicles, with rapid trafficking thought to exist between the two microdomains (7). To test whether trafficking contributes to pressure regulation of Ca_V_1.2, we implemented a stepwise set of whole-cell (rat) and single-channel (mouse) experiments in which trafficking was targeted by a microtubule disruptor (Nocodazole (1μM)), a motor protein blocker (Dynapyrazole-A (5 μM)) and a caveolae associated protein inhibitors (Caveolin-1 scaffolding domain peptide; 10μM) (**Fig. 3A, flowchart**). Whole-cell experiments revealed that each agent abolished the swelling-induced increase in Ca_V_1.2 current (**Fig. 3B-D**), with microtubule disruption additionally elevating basal activity (**Fig. 3D**). We next moved to cell-attached experiments, focusing efforts Nocodazole’s impact, as the Caveolin-1 scaffolding domain peptide and Dynapyrazole-A lack membrane permeability (**Fig. 3E**). High basal activity was evident following Nocodazole preincubation and note the ensuing inability of negative pressure to further increase single-channel activity and coupling behavior (**Fig. 3F-H**). Cumulatively, this data suggests that Ca_V_1.2 trafficking to the plasma membrane is a factor in transducing the mechano-response to pressure.

**Figure 3.**
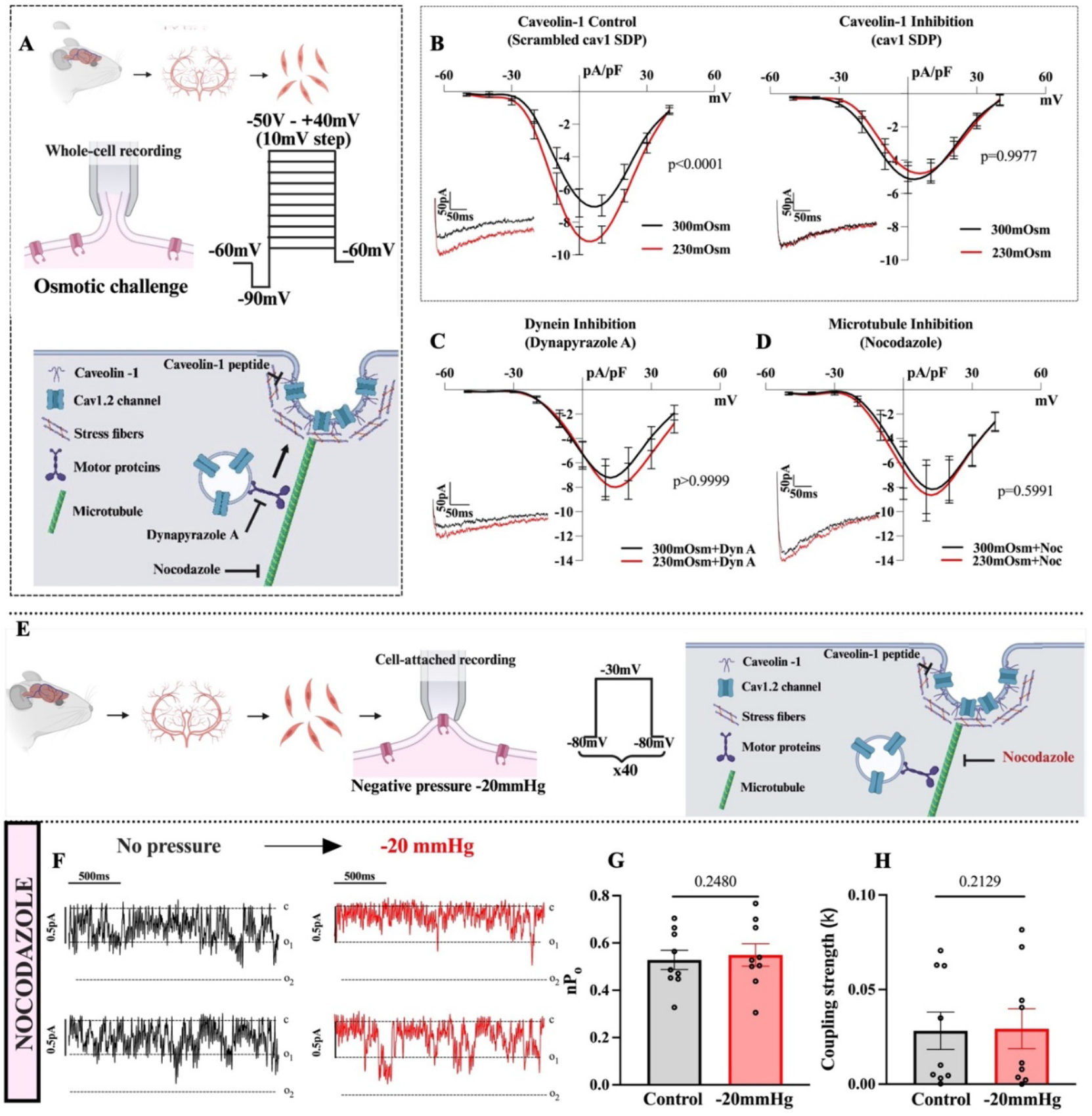
Mechano-stimulation increases whole-cell Ca_V_1.2 current by enhancing protein trafficking. **A(top)**, Schematic of cerebral arterial smooth muscle cell isolation and whole-cell patch-clamp protocol. To simulate membrane stretch by intravascular pressure, cells were exposed to a hypoosmotic bath solution (300-to-230 mOsm). **A(bottom)**, Schematic of trafficking of functionally coupled Ca_V_1.2 channels to caveolae. Microtubule dissociation, dynein inhibition, and caveolin-1-associated cytoskeletal disruption were achieved by adding Nocodazole, Dynapyrazole A, and the Caveolin-1 scaffolding domain peptide (cav1 SDP) to the pipet solution. **B**, Inhibition of caveolin-1 with cav1 SDP (10µM) abolishes the hypoosmotic-induced increase in whole-cell Ca_V_1.2 current; the control scrambled cav1 SDP (10µM) peptide has no effect (left – control, right – experiment; n=6 from 6 animals, two-way ANOVA, Sidak’s multiple comparison test, p shown for the peak value). The dynein motor protein inhibitor, Dynapyrazole A (5 µM, n=7 cells from 7 animals) (**C)**, and the microtubule disruptor, Nocodazole (1 µM, n=6 cells from 6 animals) (**D**) also impaired the hypoosmotic-induced increase in whole-cell Ca_V_1.2 current (no significant difference determined using two-way ANOVA, Sidak’s multiple comparison test, p shown for the peak value). **E**, Schematic of the cerebral arterial smooth muscle cell isolation and the cell-attached single-channel recording patch-clamp protocol. To simulate membrane stretch, negative pressure (-20 mmHg) was applied to the back of the recording pipette. **F**, Representative traces of cell-attached, single channel Ca_V_1.2 activity in the presence of the microtubule disruptor Nocodazole (1 µM), prior to and following negative pressure (-20 mmHg) application. The statistical analysis of single-channel recordings demonstrates that in cells pretreated with Nocodazole (1 uM), negative pressure application did not affect Ca_V_1.2 open probability (**G**) or coupling strength (**H**) (n=9 cells from 9 animals, Wilcoxon matched-pairs signed-rank test with a one-tailed hypothesis).

We subsequently performed microscopy experiments to provide structural evidence of the preceding and to explore the spatiotemporal events associated with mechano-stimulation. We first used the proximity ligation assay to ascertain the colocalization of Ca_V_1.2 with Caveolin-1, PKCα, and AKAP150 under iso- and hypoosmotic conditions. Protein colocalization was observed in all three scenarios, with a marked enhancement in hypoosmotic solutions **(Suppl. Figure S3-6**), suggestive though not definitive for Ca_V_1.2 subunits being trafficked to the membrane. We next performed a detailed super-resolution microscopy analysis to monitor the cluster size and density of Ca_V_1.2, AKAP150, Caveolin-1, and PKCα in smooth muscle cells. Hypoosmotic solution increased the cluster size of Ca_V_1.2, Caveolin-1, and PKCα, but had a smaller effect on AKAP150 (**Fig. 4C–F**). Ca_V_1.2 clusters showed substantial overlap with Caveolin-1, AKAP150, and PKCα under control conditions, and this overlap increased during cell swelling (**Fig. 4B**). With respect to specific cluster alignments, we found that hypoosmotic challenge markedly enhanced colocalization between Ca_V_1.2 and AKAP150 or PKCα; there was also a trend toward increased colocalization between Ca_V_1.2 and Caveolin-1 (**Fig. 4G–I**). Pretreatment of smooth muscle cells with Nocodazole or Calphostin C prevented pressure-induced Ca_V_1.2 cluster growth, consistent with Ca_V_1.2 subunit trafficking via a PKC-dependent mechanism (**Fig. 5**).

**Figure 4.**
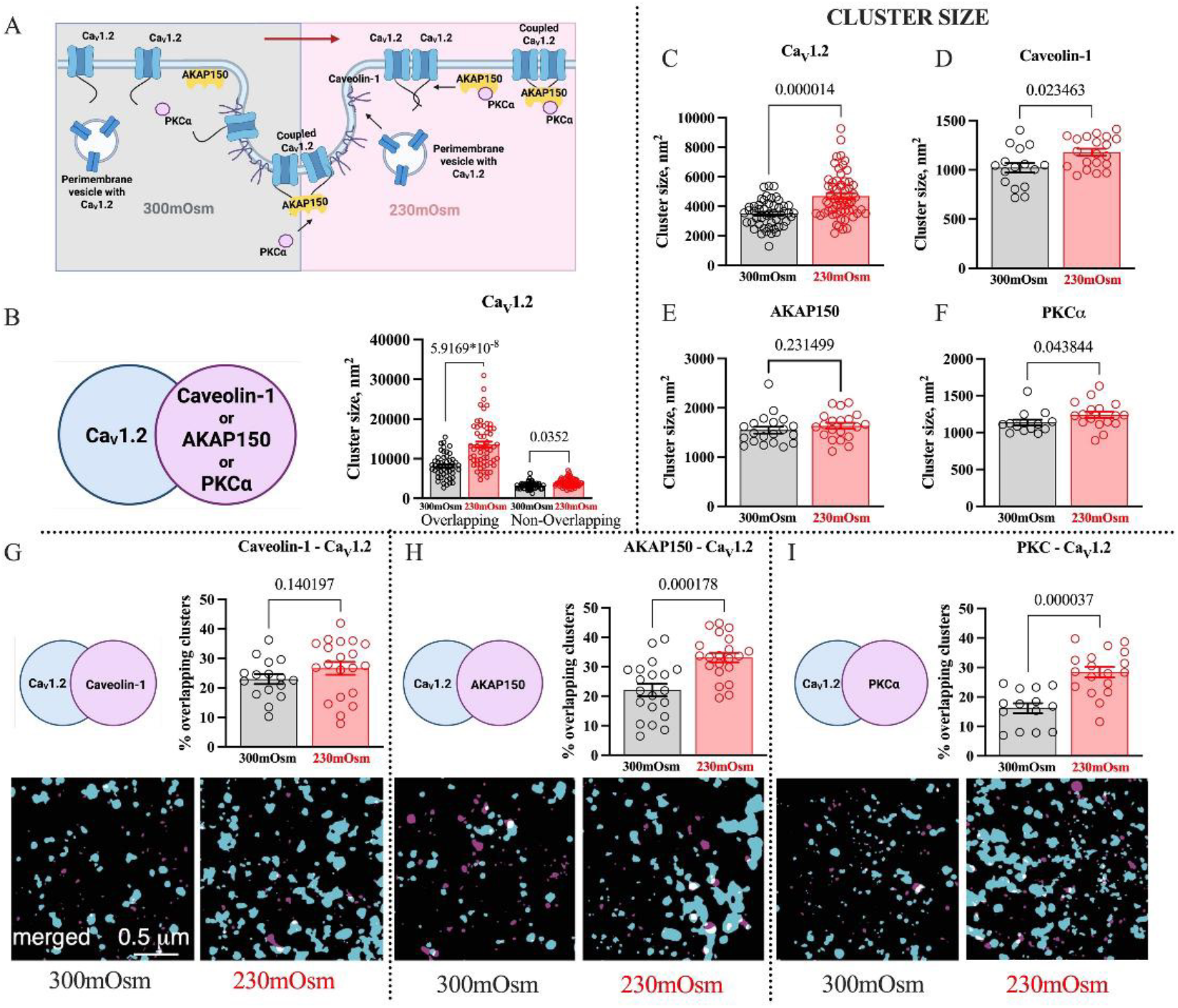
Pressure-like stimulation promotes Ca_V_1.2 cluster growth and enhanced colocalization with proteins necessary for functional coupling. **A**, Schematic panel illustrating Ca_V_1.2 trafficking dynamics in vascular smooth muscle cells in response to elevated intravascular pressure. Membrane stretch triggers fusion of perimembrane vesicles containing Ca_V_1.2 channels with the plasma membrane. Simultaneously, AKAP150 anchors PKCα to the Ca_V_1.2 C-terminus, promoting increased spatial colocalization and functional coupling. Together, these processes lead to larger Ca_V_1.2 clusters, greater channel density at the membrane, and enhanced Ca^2+^ influx. **B**, Super-resolution imaging shows that osmotic challenge, a pressure-like stimulus, significantly increases Ca_V_1.2 cluster size, particularly in clusters colocalizing with AKAP150, PKCα, or Caveolin-1 (Mann-Whitney test, n of cells =50 for 300mOsm and 59 for 230mOsm calculation, n of animals = 3). **C–F**, Quantification of protein cluster size following osmotic challenge. Ca_V_1.2 **(C)**, Caveolin-1 **(D)**, and PKCα **(F)** clusters show a significant increase in size, whereas AKAP150 **(E)** cluster size remain unchanged, (Mann-Whitney test with two tailed hypothesis, n of cells for 300mOsm and 230mOsm calculations are C: 49 and 59, D: 16 and 20, E: 20 and 20, F: 14 and 17, respectively, n of animals = 3 in each experiment). **G–I**, Osmotic challenge enhances colocalization of Ca_V_1.2 with Caveolin-1 **(G**), AKAP150 **(H)**, and PKCα **(I**). Notably, AKAP150 and PKCα—key mediators of functional coupling—exhibit significantly increased colocalization with Ca_V_1.2 under osmotic stimulation (Mann-Whitney test with two tailed hypothesis, n of cells for 300mOsm and 230mOsm calculations are G: 16 and 20, H: 20 and 21, I: 14 and 18, respectively, n of animals = 3 in each experiment).

**Figure 5.**
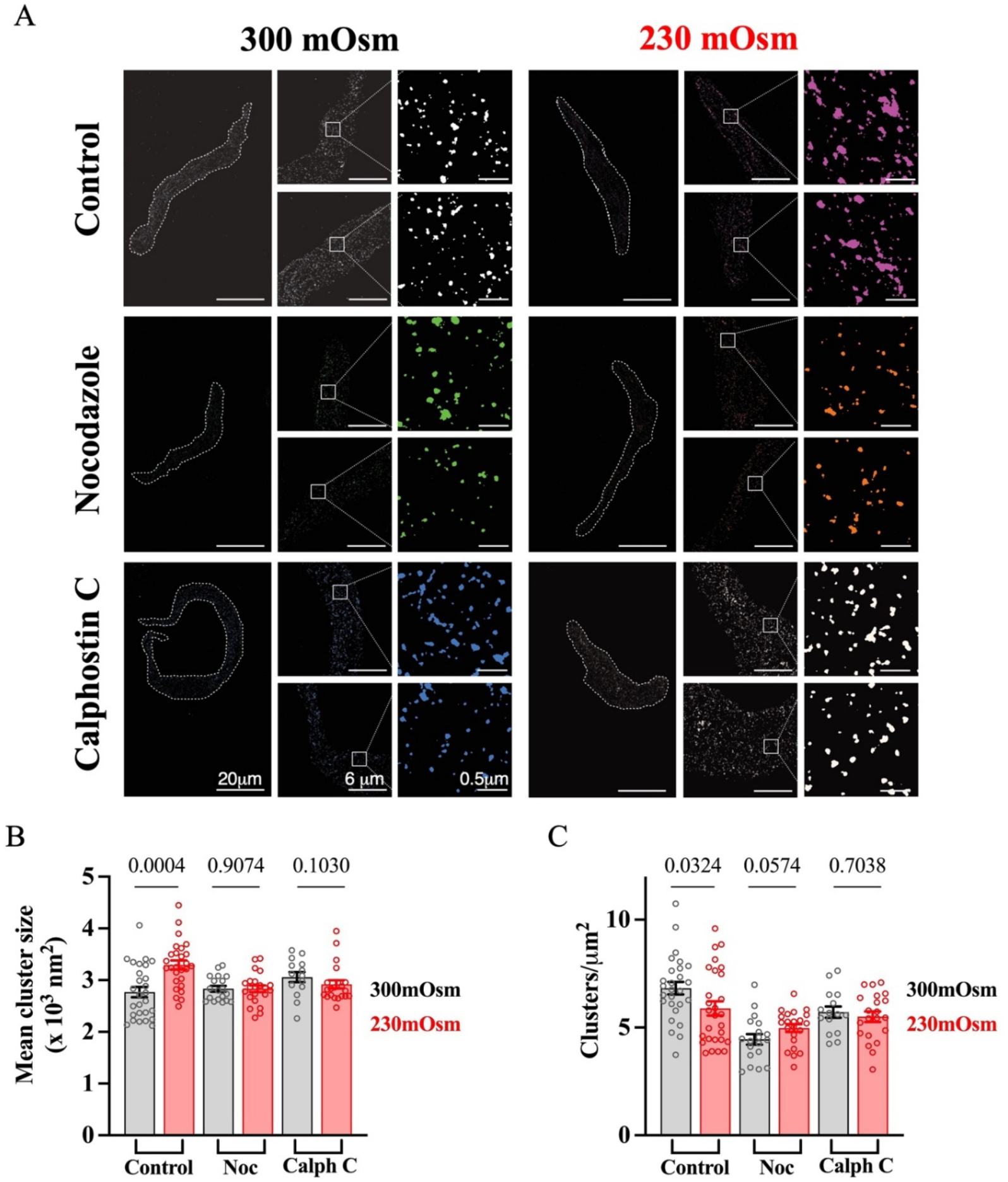
Mechano-stimulation of mouse smooth muscle cells enhances Ca_V_1.2 cluster size through protein trafficking and PKC activation. Super-resolution images (**A**) reveal the dynamic changes in Ca_V_1.2 channel clusters in smooth muscle cells exposed to a hypoosmotic challenge (230 mOsm). The top panel compares Ca_V_1.2 clusters in control cells before and after the hypoosmotic challenge; the middle and bottom panels make the same comparison between cells pretreated with Nocodazole (1 µM) or Calphostin C (300 nM), respectively. Analysis revealed the rise in Ca_V_1.2 cluster size (**B, Control**) and decrease of cluster density (**C, control**). Of note, pretreatment of cells with Nocodazole and Calphostin C inhibited the cluster size and density changes after hypoosmotic challenge (**B-C, Noc&Calph C**). Bar graphs were analyzed using Mann-Whitney test, two-tailed hypothesis, **B** (left-to-right): n of cells = 27, 27, 19, 22, 14, 21; **C** (left-to-right): n of cells = 27, 28, 19, 22, 15, 21; three animals were used per experimental group: Control, Noc, and Calph C.

### Functional Implications of Mechano-Regulated Ca_v_1.2 Channels

We next tested whether functional coupling contributes to myogenic tone, reasoning that enhanced channel cooperativity is required to counter rising wall stress with pressurization, unlike with an agonist (**Fig. 6B-E**). We consequently developed a bioassay in which arteries loaded with Fura-2AM were used to assess [Ca^2+^]_i_ response at low and high pressure (20 and 80 mmHg) in the presence of 30 mM KCl to standardize membrane potential (**Suppl. Fig.S7**). We reasoned, with membrane potential fixed, a pressure related difference in [Ca^2+^]_i_ would reflect the enhanced Ca_V_1.2 functional coupling/trafficking. Moving stepwise through these results, constriction to 30 mM KCl was similar at 20 mmHg or 80 mmHg (**Fig. 6G**), while [Ca^2+^]_i_ was higher at 80 mmHg (**Fig. 6I**). These findings align with enhanced functional coupling/trafficking being essential for greater force generation needed counter to elevated wall stress with pressurization. Interference with PKC activity (Calphostin C/Go 6976) or perimembrane trafficking (Nocodazole) moderated the rise in [Ca^2+^]_i_ at 80 mmHg (**Fig. 6H**). We next used the differential [Ca^2+^]_i_ response, 1) 20 mmHg+30 mM KCl - 20 mmHg and 2) 80 mmHg+30 mM KCl - 20 mmHg+30 mM KCl, as a measure of Ca^2+^ channel activity tied to V_M_ and functional coupling. The plot of relative contribution (**Fig. 6J**). reveals that Vm and functional coupling are near equally important to driving the pressure-induced [Ca^2+^]_i_ response.

**Figure 6.**
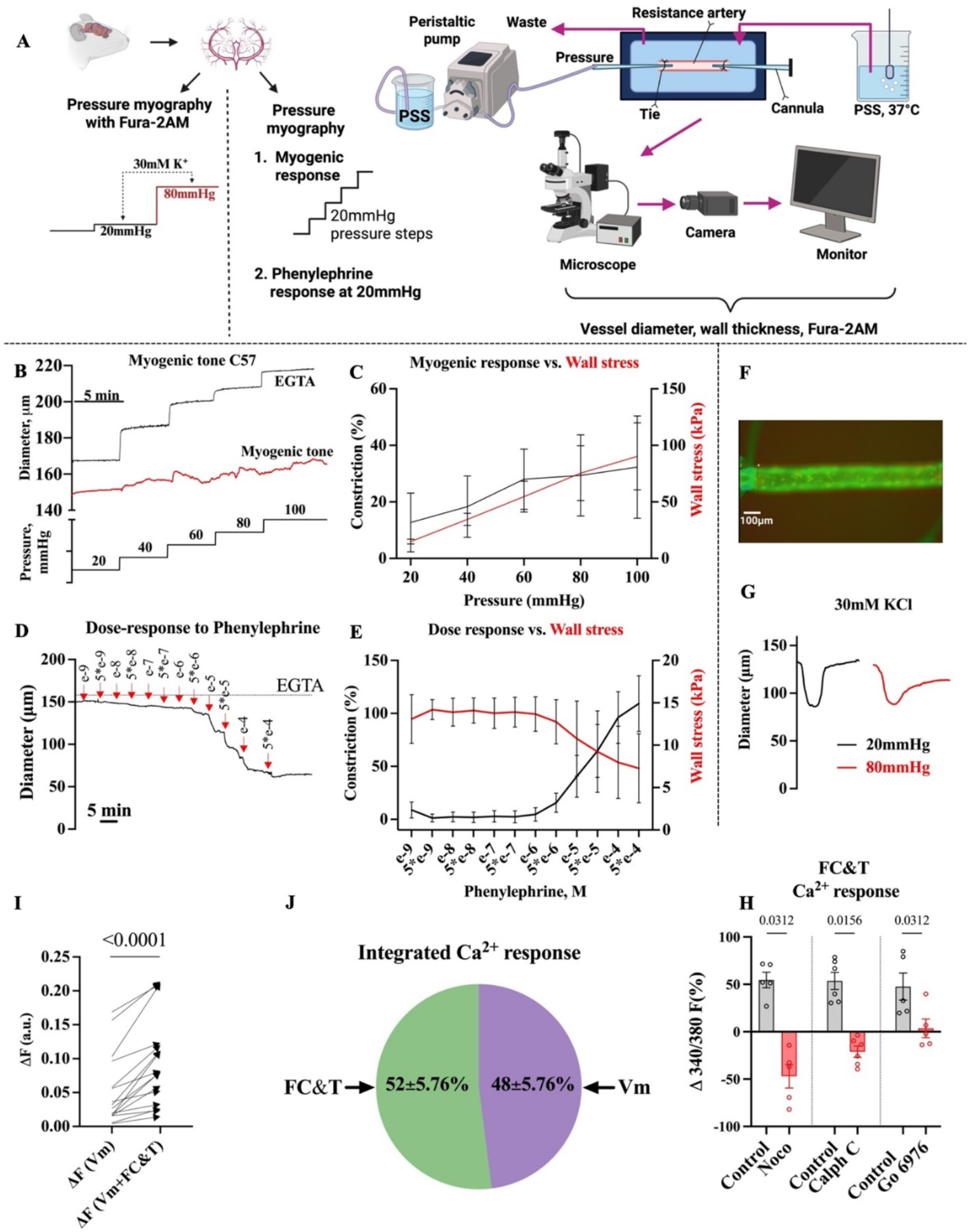
Myographic bioassay of mouse cerebral arteries reveals that functional coupling increases with pressurization. **A**, Schematic of mouse brain arteries isolation, and the two experimental protocols. For pressure myography assay cerebral arteries were mounted in a customized pressure myograph and loaded with Fura-2AM to monitor [Ca^2+^]_i_. Vessels were held at low (20 mmHg) and high (80 mmHg) pressure while diameter and [Ca^2+^]_i_ were monitored under control conditions and after the addition of 30 mM extracellular K^+^ to elicit a standardized voltage response to activate Ca_V_1.2 channels. **B**, Representative trace myogenic tone development in response to increased intravascular pressure; **C**, Summary highlights the relationship between myogenic constriction and wall stress (n=6). **D**, Representative trace of arterial constriction to increasing phenylephrine concentration; **E**, Summary highlights the relationship between phenylephrine-induced constriction and wall stress (n=6). **F**, Representative image of a cerebral artery mounted in the arteriograph and loaded with Fura-2 AM. **G**, Representative constrictor trace of an artery (set at 20 and 80 mmHg) to 30 mM KCl. **H**, Percentage of ΔF attributed to trafficking/functional coupling in control vessels and those pretreated with nocodazole (Noco, 10 µM), Calphostin C (Calph C, 300 nM), or Go 6976 (Go, 100 nM) at 80 mmHg. Data analyzed using one-tailed Wilcoxon matched-pairs signed-rank test (Noco: 5 vessels from 5 mice, Calph C: 6 vessels from 6 mice: Go: 6 vessels from 6 mice). **J**, Sector diagram illustrating the relative contributions of membrane depolarization (Vm, purple) and functional coupling with trafficking (FC&T, green) to the total change in intracellular Ca^2+^ fluorescence (ΔF) in mouse cerebral arteries in response to 30 mM KCl constriction at 80 mmHg. Values are presented as mean ± SEM (n = 16 vessels from 16 mice). **I**, Changes of [Ca^2+^]_i_ fluorescence (vessels set at 20 and 80 mmHg) to 30mM KCl (n=16 vessels from 16 mice, Wilcoxon matched-pairs signed-rank test with a one-tailed hypothesis).

### Genetic inference with Ca_V_1.2 functional coupling

To further probe the link between functional coupling and myogenic tone, myographic experiments were conducted on arteries from Ca_V_1.2-S1928A mice, a mutation that interferes with protein kinase phosphorylation (16). Compared to age-matched C57BL/6 controls, cerebral arteries from S1928A mice (**Fig. 7A; Fig. 7C–D, illustrative traces**) generally displayed reduced tone, with pressure-induced responses preserved up to 60 mmHg, consistent with a maintenance of voltage-dependent regulation. Above 60 mmHg, transgenic arteries failed to generate the additional myogenic tone needed to counter increased wall stress (**Fig. 7E–F**). Note cell-attached patch-clamp experiments confirmed reduced Ca_V_1.2 open probability/functional coupling under basal conditions, and the absence of pressure-induced enhancement in cerebral arterial smooth muscle from S1928A transgenic mice (**Fig. 7G–L**).

**Figure 7.**
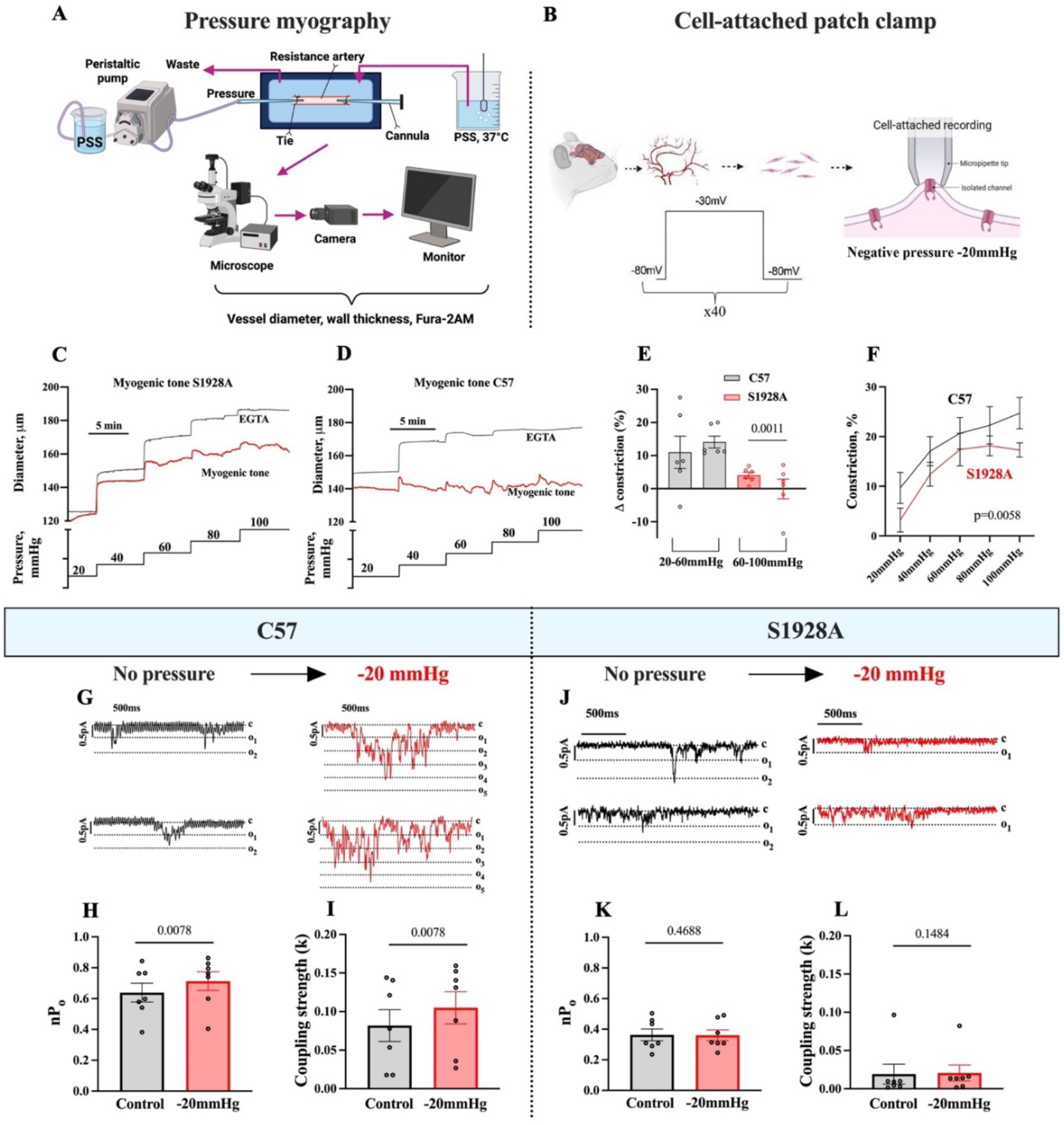
Pressure myography and patch-clamp electrophysiology reveals phosphorylation-dependent regulation of Ca_V_1.2 channel activity and myogenic response in mouse cerebral arteries. **A**, Schematic of the pressure myography setup for isolated pressurized cerebral arteries. Arteries were cannulated and pressurized in physiological saline solution (PSS) at 37°C, and vessel diameter was continuously monitored using video microscopy. **B**, Schematic of the cell-attached patch-clamp configuration on isolated cerebral artery smooth muscle cells. To simulate mechanical stress, negative pipette pressure (-20 mmHg) was applied. **C-D**, Representative traces showing myogenic tone development over time in arteries from S1928A mice and C57 controls, respectively. **E**, Quantification of pressure-induced constriction in S1928A arteries across two pressure ranges (20–60 mmHg and 60–100 mmHg) using a one-tailed Mann–Whitney test (nonparametric, unpaired), indicating a directional reduction in tone at higher pressures (n = 6 vessels from 6 animals). **F**, Summary data showing pressure-dependent myogenic constriction in C57 and S1928A arteries (n = 6 cells from 6 animals; two-way ANOVA, Sidak’s multiple comparisons). **G-J**, Representative Ca_V_1.2 single-channel currents recorded from C57 (**G**) and S1928A (**J**) smooth muscle cells under control conditions and following application of -20 mmHg pipette pressure. **H-I**, Summary of open probability (nPo) and coupling strength (k) for Ca_V_1.2 channels in C57 cells, showing significant increases upon -20 mmHg stimulation (n = 7 cells from 7 animals, Wilcoxon matched-pairs signed-rank test with a one-tailed hypothesis). **K-L**, In contrast, S1928A cells showed no significant change in nPo or coupling strength under the same conditions (n = 6 cells from 6 animals, Wilcoxon matched-pairs signed-rank test with a one-tailed hypothesis).

### Mechano-Stimulation of Ca_V_1.2 Channels in Human Cerebral Arteries

We repeated key experiments using human cerebral arteries obtained from resected brain tissue of epileptic patients (**Fig. 8A flowchart**). Similar to mice, application of negative pressure (–20 mmHg) via the patch pipette increased single-channel Ca_V_1.2 activity (**Fig. 8B)**, and quantitative analysis confirmed that pressure elevated both open probability and coupling strength (**Fig. 8E– F**). In further alignment, we observed when membrane potential was equalized with 30 mM KCl that [Ca^2+^]_i_ was higher at 80 mmHg compared to 20 mmHg (**Fig. 8C**). This finding aligns with functional coupling contributing substantive to the myogenic Ca^2+^ response, an interpretation best observed in the relative contribution plot (**Fig. 8D**).

**Figure 8.**
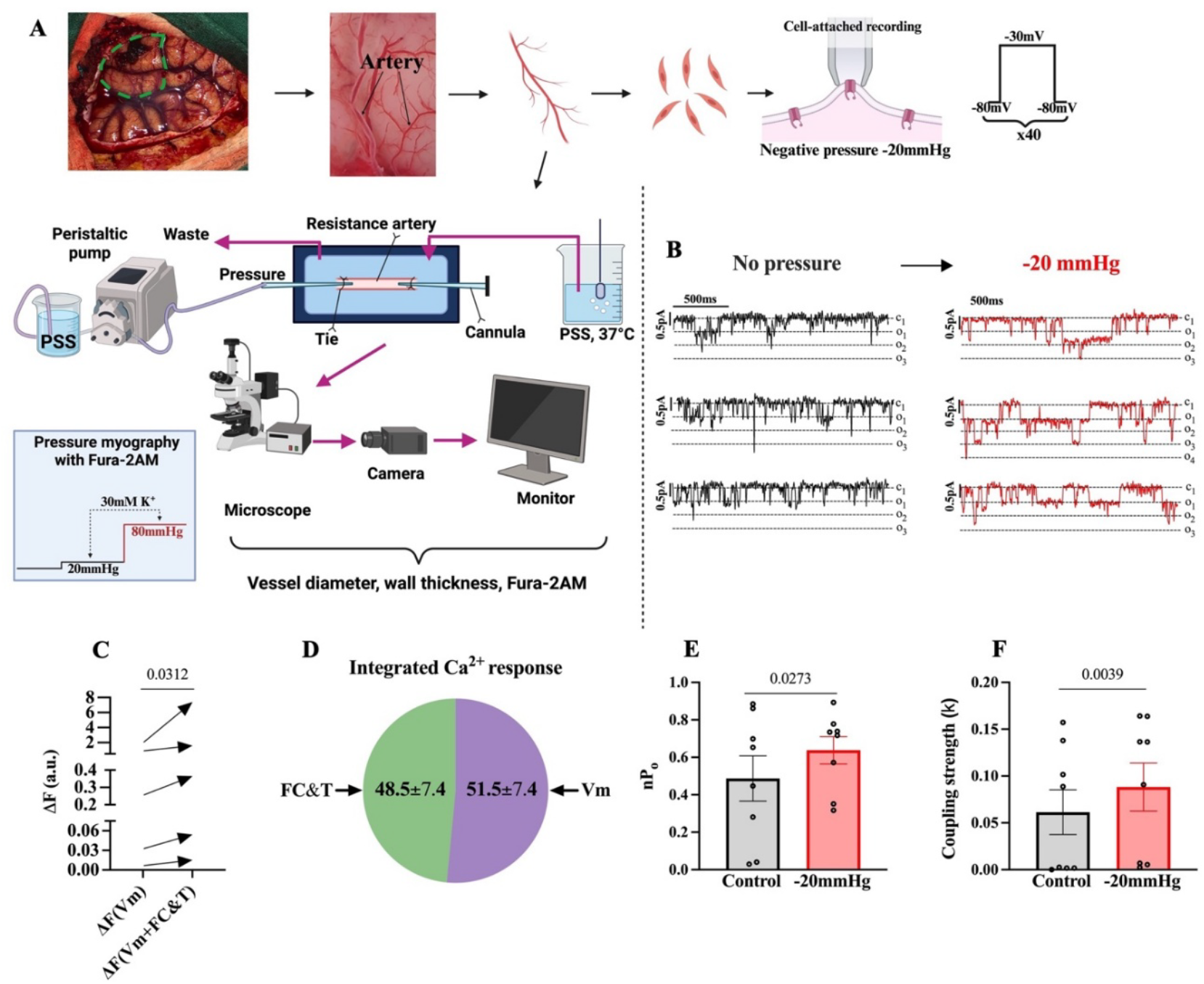
Mechano-stimulation increases Ca_V_1.2 single-channel activity in human cerebral arterial smooth muscle cells. **A**, Schematic of human brain resection surgery, arteries isolation, and the two experimental protocols. For pressure myography assay cerebral arteries were mounted in a customized pressure myograph and loaded with Fura-2AM to monitor [Ca^2+^]_i_. Vessels were held at low (20 mmHg) and high (80 mmHg) pressure while diameter and [Ca^2+^]_i_ were monitored under control conditions and after the addition of 30 mM extracellular K^+^ to elicit a standardized voltage response to activate Ca_V_1.2 channels. In cell-attached patch-clamp assay, to simulate membrane stretch, negative pressure (-20 mmHg) was applied via the pipet on isolated arterial smooth muscle cells. **B**, Representative traces of Ca_V_1.2 single-channel activity before and after negative pipet pressure application (-20 mmHg). Analysis demonstrated that negative pressure application increased open probability (**E**) and coupling strength (**F**) of Ca_V_1.2 channels (n=8 cells from 7 humans, Wilcoxon matched-pairs signed-rank test with a one-tailed hypothesis). **C**, Changes of [Ca^2+^]_i_ fluorescence (vessels set at 20 and 80 mmHg) to 30mM KCl (n=5 vessels from 5 humans, Wilcoxon matched-pairs signed-rank test with a one-tailed hypothesis). **D**, Sector diagram illustrating the relative contributions of membrane depolarization (↑Vm, purple) and functional coupling with trafficking (green) to the total change in intracellular calcium fluorescence (ΔF) in a human vessel in response to 30 mM KCl constriction at 80 mmHg. Values are presented as mean ± SEM (n = 5 vessels from 5 humans).

### In silico experiments: Impact of functional coupling on blood flow

Using a semi-realistic microvascular network model comprised of surface arteries, penetrating arterioles, a capillary bed and ascending veins (**Fig 9A**, (17)), we probed how brain blood flow distribution would likely change in S1928A mice. The S1928A phenotype was simulated by reducing the myogenic sensitivity factor (*S*_*σ*_=4.0) by 30% (**Fig. 9B flowchart**), in line our data collected at 100 mmHg (**Fig. 7F**). We first noted an attenuated constrictor response in arteries that we stratified according to intraluminal pressure (**Fig. 9C-9D**: surface arteries (80-100 mmHg) vs penetrating/pre-capillary arterioles (20-40 mmHg). This attenuation had marked impact on the ensuing blood flow response induced by transiently elevating inlet pressure in our model. Note the maldistribution of relative capillary flow, with regions on the surface being oversupplied while those deeper (**Fig. 9E**, 600-800 μm) were undersupplied.

**Figure 9.**
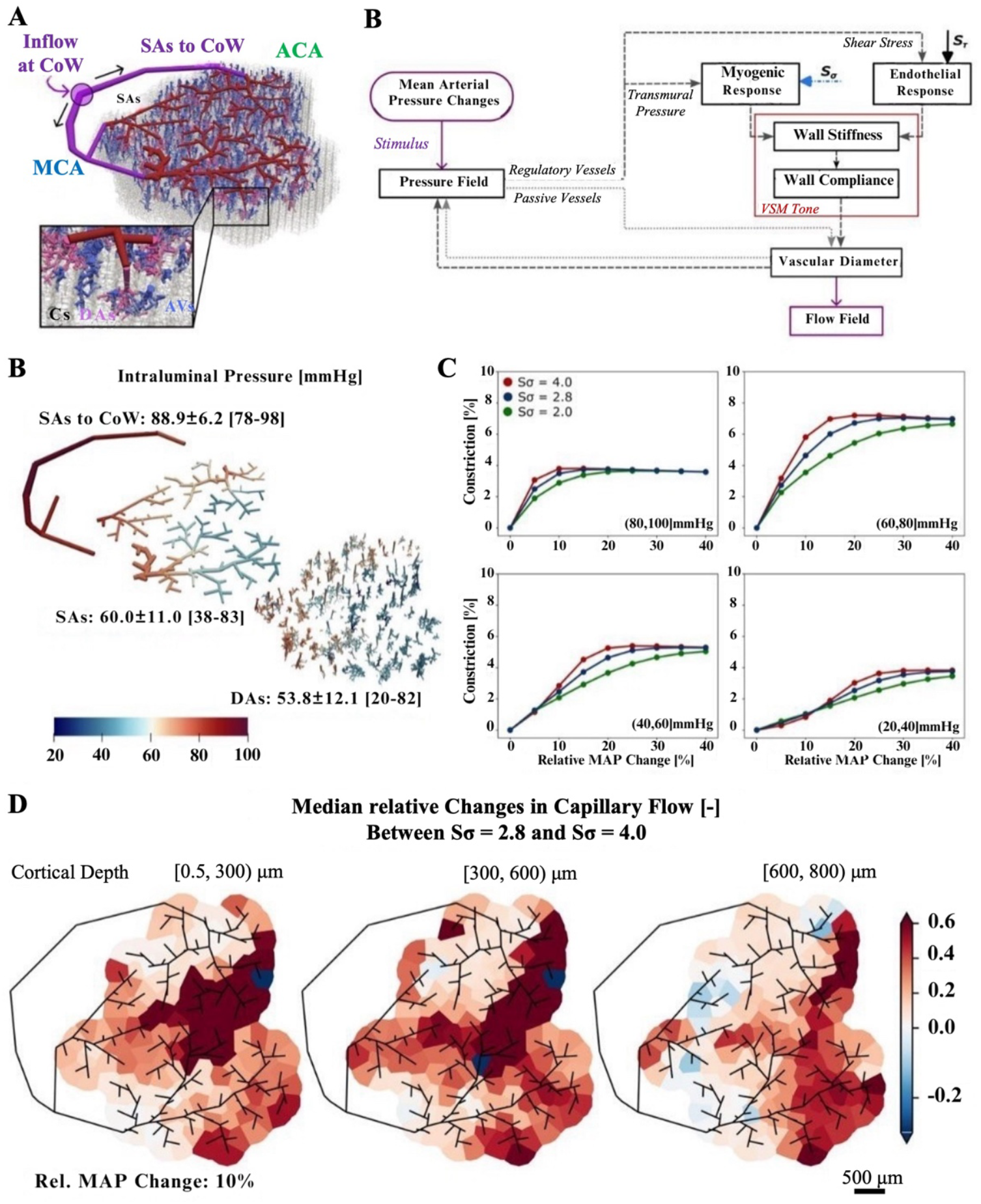
Impact of myogenic impairment on arterial constriction and capillary perfusion under elevated pressure. **(A)** Semi-realistic microvascular network (MVN) based on the realistic surface arterial (SA) topology of a Balb-C mouse. The zoom-in shows the added penetrating trees (descending arterioles, DAs, and ascending venules, AVs) and the artificial capillary bed (Cs). **(B)** Schematic representation of the autoregulation model. The flow chart for active and passive vessels is depicted by dashed and dotted lines, respectively. **(C)** Intraluminal pressure (IP) distribution across all arterial segments at baseline conditions. SAs-to-CoW: surface arteries connecting to the Circle of Willis. **(D)** Arterial constriction as a function of relative mean arterial pressure (MAP) change for three different myogenic sensitivity factors (*Sσ*=4.0, 2.8, 2.0). Arteries are grouped based on their baseline intraluminal pressure (IP): the left panels show results for IP ∈ (60, 80] mmHg, and the right panels, IP ∈ (40, 60] mmHg. **(E)** Spatial distribution of changes in capillary perfusion due to reduced myogenic sensitivity for a 10% increase in MAP. Capillaries are grouped in columns below each DA root point to allow a planar representation. Each panel shows the median relative difference in capillary flow between *Sσ* = 2.8 and *Sσ* = 4.0, computed as *median*(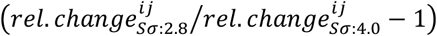) across all capillaries in a given cortical depth range: [0.5, 300), [300, 600), and [600, 800] μm. Red regions indicate an increase in flow under impaired myogenic response, with more pronounced effects observed in superficial layers.

## Discussion

The myogenic response is foundational to blood flow regulation and prototypically viewed as an autoregulatory constriction to elevated pressure (1, 2, 18). The canonical mechanism entails pressure eliciting an arterial depolarization that sequentially increases [Ca^2+^]_i_ and myosin light chain phosphorylation in support of actin–myosin cross-bridge formation (19–21). The voltage dependent rise in [Ca^2+^]_i_ is tied to the activation of L-type Ca^2+^ channels, a multimeric complex comprised of an α1 pore (Ca_V_1.2) and auxiliary subunits (α2, β, *δ*, and γ) that control gating, anchorage, and trafficking. Each Ca_V_1.2 subunit retains four homologous domains, each of six transmembrane segments, along with an N- and C-termini - the latter a key site of regulation. The S4 segment is the “voltage sensor” and upon depolarization, it moves relative to S5-S6 to open the pore (3, 4). As pressure notably depolarizes smooth muscle, voltage is considered the primary if not sole factor driving Ca_V_1.2 activation. But consider the autoregulatory window of cerebral arteries (50 to 150 mmHg) relative to the voltage window (-35 to -50 mV), and if the latter alone can provide the precision tuning for the former. Is there in essence an additional regulatory mechanism by which pressure regulates L-type Ca^2+^ channels? This idea drives our interest in functional coupling, a process whereby Ca_V_1.2 subunits interact via their C-termini to increase open probability. Functional coupling is intimately tied to the clustering of Ca_V_1.2 subunits, a process controlled by key kinases (PKC and PKA) tethered to Ca_V_1.2 C-termini and potentially to the trafficking of coupled Ca_V_1.2 subunits from the perimembrane space (6, 7). While functional coupling has received noted attention in relation to pathology (8–10), its role in general physiology has been overlooked - an issue we address herein.

### Electrophysiology: Mechano-Stimuli and Functional Coupling

The work of Knot & Nelson (1998) was seminal in outlining a mechanistic framework to rationalize how pressure enhances cerebral arterial tone (22). That framework entailed pressure inducing depolarization to trigger L-type Ca^2+^ channels and elevate [Ca^2+^]_i_ so to activate myosin light chain kinase. Interest in the ion channels underlying depolarization ensued, key candidates being transient receptor potential, and inward rectifying K^+^, channels (23). The effector itself, L-type Ca^2+^ channels, received markedly little attention as to its pressure sensitivity, outside a small subset of rat studies (11–15). Aligned with this unacknowledged work, cell swelling notably increased the whole-cell Ca_V_1.2 current, a stimulus that stretches the plasma membrane and which is used to identify pressure-regulated channels (**Fig. 1**) (24). Calphostin C, a pan PKC inhibitor, attenuated this swelling-induced response, an observation that indirectly suggests that enhanced functional coupling underlies the whole-cell response. Ensuing cell-attached measures, conducted on mouse rather than rat cerebral arterial smooth muscles cells confirmed that pipette pressure application increases single channel Ca_V_1.2 activity, including events where two or more channel opened together (**Fig. 2)**. The latter reflected in nPo and coupling strength are standard indices of functional coupling and note these changes were absent in cells pretreated with Calphostin C. Within the PKC family, PKCα is known to govern L-type Ca^2+^ channel cooperatively (25)(26, 27)Go6976, a selective PKCα inhibitor abolished these pressure-induced responses. Deeper examination revealed that both PKC inhibitors elevated resting single channel activity including coupled events, a result difficult to solely rationalize through PKC phosphorylating the Ca_V_1.2 C-terminus. In this regard, this kinase could be controlling a broader set of cellular events, the most apparent being subunit trafficking to the plasma membrane (7, 28).

### Electrophysiology: Mechano-Stimuli, Trafficking and Functional Coupling

In cultured cell systems, Ca_V_1.2 subunits traffic between the plasma membrane and the perimembrane space (7, 29), a process guided along microtubule tracts by motor proteins like dynein. They “walk” vesicles to caveolae where they merge with existing clusters (30, 31). As interference of Ca_V_1.2 trafficking has been previously shown to attenuate tone (7, 32), we deduced it might also contribute to the transduction of pressure-induced functional coupling. This perspective aligned with our initial whole-cell measures (rat cerebral arterial smooth muscle) where microtubule disruption, dynein inhibition and Caveolin-1 binding interference each attenuated the swelling-induced response (**Fig. 3**). Note our dynein focus was driven by inhibitor availability and its purported role in trafficking preexisting vesicles from perimembrane pools rather than those newly synthesized (7, 30, 33). More direct linkage was observant in single-channel work where Nocodazole preincubation (membrane permeable microtubule disruptor) blocked pressure-induced functional coupling in mouse cerebral arterial smooth muscle. Microtubule disruption also elevated basal channel activity, a mirror of our PKC work, and perhaps indicative of this kinase’s role in pressure-induced trafficking (34)

In considering the preceding work, two structural relationships become apparent, each requiring careful verification. The first centers on Ca_V_1.2 co-localizing with AKAP and PKCα within key mechano-structures (e.g. caveolae), relationships first confirmed by proximity ligation assay, which detects two proteins residing within 40 nm of one another. These focal red puncta evident in isosmotic solutions saw their formation rise in hypoosmotic solutions, consistent but not definitive for, pressure stimulation of Ca_V_1.2 trafficking (**Suppl. Figure S3-6**). More precise analysis enabled by super-resolution microscopy (**Fig. 4**), showed how hypoosmotic stress also increased Ca_V_1.2 cluster size as it did other key trafficking/coupling proteins (Caveolin-1 and PKCα), supportive of coordinated regulation amongst Ca_V_1.2 with it signaling machinery. Deeper analysis revealed that Ca_V_1.2 clusters also overlapped with trafficking and coupling proteins, an association that rose with pressure-like stimulation. These pressure-induced responses were abolished following Nocodazole and Calphostin C preincubation, the latter supportive of PKC being involved in the channel trafficking process **(Fig. 5)**.

### Vessel Myography and Functional Coupling/Trafficking

Constriction is the principal vasomotor event in the cerebral circulation as without it functional hyperemia ceases and there is no effective means to set blood flow distribution or capillary pressure. Pressure is the key stimulus in setting base tone, a response exclusively tied to voltage control of L-type Ca^2+^ channels. While depolarization is critical, we queried whether mechano-stimuli had an additional regulatory on L-type Ca^2+^ channels via functional coupling to enhance the influx of Ca^2+^ needed to counter the rise in wall stress (**Fig. 6**). To test whether pressure enhanced functional coupling *in situ*, we implemented a custom bioassay in which isolated cerebral arteries were pressurized to 20 or 80 mmHg, then briefly exposed to 30mM K^+^ to standardize smooth muscle Vm (**Suppl. Figure S7**). We rationalized that if pressure enhanced functional coupling, [Ca^2+^]_i_ would be greater at 80 rather than 20 mmHg. This was observed and note the equal partition of the Ca^2+^ response to that of voltage control or functional coupling, the latter blocked by Calphostin C, Go-6976, and Nocodazole. One can conclude from this vessel-level work that functional coupling is indeed essential for myogenic tone generation and that PKCa likely mediates this through 1) phosphorylation of Ca_V_1.2 C-terminus, and/or 2) augmentation of subunit trafficking from the perimembrane space. The latter aligns with the work of Jaggar and colleagues who showed that surface expression of Ca_V_1.2 is indeed pressure dependent in this tissue (35).

### Genetic Impairment of Ca_V_1.2 Phosphorylation

To further extend our knowledge of pressure-induced functional coupling, we examined mice carrying the Ca_V_1.2-S1928A substitution, a mutation that impairs kinase phosphorylation and which knowingly impacts channel cooperativity (36). This mutation reduced overall tone in a unique manner, with pressure responsiveness maintained between 20 and 60 mmHg but muted between 60 and 100 mmHg (**Fig. 7**). These results suggest a sequential arrangement of mechanisms, with depolarization driving the initial activation of Ca_V_1.2, followed by functional coupling key at higher pressures where Vm is near its functional maximum. Note prudent single channel recordings confirmed pressure-induced functional coupling was indeed absent in smooth muscle cells from this transgenic line.

### Human Integration

Animal models serve as a valuable test bed to explore foundational concepts and to set a platform for human translation. In this regard, we procured fresh human brain tissue and harvested surface arteries with the goal of validating key experiments. With patch-clamp electrophysiology, L-type Ca^2+^ channels were identified at the single channel level and their marked sensitivity to negative pressure application documented, including a rise in functional coupling as denoted by the change in nPo and coupling strength (**Fig. 8B, E-F**). In alignment, myography work confirmed that like mouse, [Ca^2+^]_i_ was higher at 80 than 20 mmHg, under conditions in which membrane potential was standardized with 30mM K^+^ (**Fig. 8C**). So again, intravascular pressure not only regulates functional coupling but facilitates myogenic tone development. This work was conducted on males and females under age 40, thus it’s unclear what happens to functional coupling in older adults. Perhaps it decreases in line with the known age-induced decline in the whole-cell current (37).

### Functional Impact and Conclusions

Computational models provide a means to integrate experimental work into an understanding of network-level behavior. In this regard, we employed a semi-realistic microvascular network with encoded myogenic and endothelial responses to probe changes in capillary blood that are likely occurring in the S1928A transgenic mouse which show reduced myogenic sensitivity (∼30%). This in silico work predicted a profound maldistribution likely occurs in this transgenic mouse, with surface regions nearest the pial arteries receiving too much blood flow whereas deeper tissue receives too little (**Fig. 9**). This behavior mirrors that of the so called “steal” phenomenon, a state known to foster microbleeds and hypoxia in regions of over and under supply, respectively (29). In summary, this study expands our mechanistic understanding of L-type Ca^2+^ channels and how they drive the key autoregulatory reflex in brain vasculature. Using a stepwise approach, we showed that in addition to voltage dependent regulation, pressure enhances L-type Ca^2+^ channels, pressure through the induction of functional coupling. This enhancement was shown to be under the regulatory control of PKCα, a kinase that facilitates cooperativity through (1) channel phosphorylation and (2) the promotion of subunit trafficking to the plasma membrane. Collectively, these findings redefine our understanding of cerebral blood flow regulation, revealing that functional coupling is a dominant PKCα-regulated driver of vascular tone and a potential nexus for vascular dysfunction and advancement of neurodegenerative disease.

## Methods

### Animal and Human Tissues

All animal procedures (except knock-in S1928A mice) were approved by Western University Animal Care Committee in accordance with the Canadian Council on Animal Care and in alignment with ARRIVE principles. This study used female Sprague-Dawley rats (10-12 weeks of age) and male C57BL mice (16-20 weeks of age), both euthanized by CO_2_ asphyxiation. Following euthanasia, the brain was removed and immediately placed in cooled phosphate-buffered solution (PBS) (pH 7.4) containing (in mM): 138 NaCl, 3 KCl, 2 NaH_2_PO_4_, 10 Na_2_HPO_4_, 5 glucose, 0.1 MgSO_4_ and 0.1 CaCl_2_. Cerebellar, middle and posterior cerebral arteries were then dissected, cleaned, and cut into 2-4 mm segments for pressure myography or enzymatic digestion (patch-clamp electrophysiology or immunohistochemistry). Arteries from female rats were initially used for two reasons: 1) they yield high quality smooth muscle cells ideal for extended whole-cell patch-clamp experiments; 2) they match past publications, thus, facilitating interstudy comparison (11, 14). Once whole-cell experiments were completed, all further tests employed mouse cerebral arteries, as it’s today’s preferred standard.

Use of S1928A transgenic mice was governed according to the Institutional Animal Care and Use Committee at the University of California, Davis. Briefly age-matched 8-10 weeks-old C57BL/6J (WT) males (Jackson Laboratory) housed in a temperature-controlled environment (21 ± 2° C, 12h light/dark photocycle) were backcrossed with females carrying the Ca_V_1.2-S1928A to prevent α1C subunit phosphorylation (16) Heterozygote breeding was used to create homozygote progeny and they in turn back breed for 10 generations. Experimental procedures including tissue isolation, solutions, and test protocols were constant amongst the groups; mice were fed food and water *ad libitum*.

Excised human brain samples were obtained after institutional review at Western University and with informed consent of the patient, and in accordance with the Declaration of Helsinki. Brain tissues were collected and immediately placed in cold PBS (pH 7.4) containing (mM) 138 NaCl, 3 KCl, 10 Na_2_HPO_4_, 2 NaH_2_PO_4_, 5 glucose, 0.1 CaCl_2_, and 0.1 MgSO_4_, and transferred to the laboratory within 10 minutes. Small superficial cerebral arteries (∼150–250-µm diameter) were carefully dissected out of surrounding tissue and cut into segments for further scrutiny same day. None of the donors underwent invasive EEG monitoring prior to tissue collection. The samples were obtained from both male and female donors, all under 40 years of age.

### Isolation of Arterial Smooth Muscle Cells

Dissected arteries were placed in an isolation media (10 min) containing (in mM): 80 sodium glutamate, 5 KCl, 60 NaCl, 2 MgCl_2_, 10 glucose, 10 HEPES and 1 mg/mL of bovine serum albumin (pH 7.4). Vessels were then settled on ice (10 min) before warming to 37 ºC for a two-step digestion process: 1) 13–15-minute incubation in media containing 0.5 mg/mL of papain and 1.5 mg/mL of dithiothreitol, and 2) 10-minute incubation in media containing 100 μM CaCl_2_, 0.3 mg/mL H-type collagenase, and 0.6 mg/mL F-type collagenase. Following enzymatic digestion, vessels were washed with ice-cold isolation media and gently triturated with a fire-polished Pasteur pipette; dissociated cells were used same day.

### Whole-Cell Patch-Clamp Electrophysiology

Whole-cell patch-clamp electrophysiology was first used to measure composite Ca_V_1.2 channels activity in cerebral arterial smooth muscle cells. Recording electrodes pulled from borosilicate glass (Sutter Instruments, Novato CA) on a micropipette puller (Narishige PP-830, Tokyo, Japan) were fire-polished (Narishige MF-830, Tokyo, Japan) and then gently placed onto a cell where negative pressure was applied to induce membrane rupture. Currents were recorded using an Axopatch 200B amplifier (Molecular Devices, Sunnydale CA) and processed with Clampfit 10.6 software (Molecular Devices, Sunnydale CA). Cells were first held in isoosmotic (300mOsm, calculated based on chemical composition) bath solution containing (in mM): 70 NaCl, 1 CsCl, 10 BaCl_2_, 1.2 MgCl_2_, 10 HEPES, 10 glucose, and 80 Mannitol (pH 7.4) to collect control data. Next, mechano-stimulation was induced by exposing cells to a hypoosmotic bath solution (230mOsm, calculated based on chemical composition) containing (in mM): 70 NaCl, 1 CsCl, 10 BaCl_2_, 1.2 MgCl_2_, 10 HEPES, and 10 glucose (pH 7.4). The pipette solution contained (in mM): 135 CsCl, 5 Mg-ATP, 10 HEPES, 10 EGTA (pH 7.2). The whole-cell voltage protocol consisted of holding at -60 mV (300 ms), pre-pulsing to -90 mV (200 ms) and then voltage stepping from -50 mV to +40 mV (300 ms, 10 mV increments). Cells were then returned to their holding potential and the protocol repeated every minute throughout the experiment. Agents used to modulate whole-cell channel activity were added to pipette or bath solutions: Nocodazole (1 µM, pipette), Dynapyrazole A (5 µM, pipette), st-Ht31 (10 µM, pipette) or its control peptide st-Ht31-P (10 µM, pipette), caveolin-1 scaffolding domain peptide (cav1 SDP; 10 µM, pipette) or a scrambled cav1 SDP (10 µM, pipette), Calphostin C (300 nM, bath). Offset potentials were minimized by placing a 1 M NaCl-agar salt bridge between the reference electrode and the bath. All recordings were performed at room temperature (22 °C).

### Single-Channel Patch-Clamp Electrophysiology

Single-channel Ca_V_1.2 activity was assessed in cell-attached mode using Ca^2+^ as a charge carrier. The bath solution contained (in mM): 145 KCl, 2 MgCl_2_, 0.1 CaCl_2_, 10 HEPES, and 10 glucose (pH 7.4) whereas the pipette solution consisted of (in mM) 20 CaCl_2_ and 10 HEPES (pH 7.2); the latter was also supplemented with S-(-)-BayK-8644 (500 nM) to promote longer open times while preserving native conditions, as established by Hess et al. (38). The voltage protocol consisted of 40 sweeps with voltage stepped from -80 mV (150ms, holding potential) to -30 mV (5000ms), and then return to -80mV (400ms). This protocol was conducted under control conditions and then with negative pipette pressure (-20 mmHg) applied on the same cell. All single-channel currents were recorded at 10 kHz and subsequently low pass filtered at 5 kHz; ∼80 sweeps were analyzed per experiment. Agents used to modulate channel activity were added to the pipette and bath solutions: Go 6976 (100 nM), Nifedipine (200 nM), Calphostin C (300 nM), Nocodazole (1 µM). The ensuing data stream was then processed with an in-house developed algorithm to obtain the number of channels in the recording (N), open probability (P_o_, nP_o_), coupling frequency (number of sweeps with coupled opening expressed relative to the total number of sweeps collected), coupling strength (κ, coupling coefficient). See **Computational Analysis** for details.

### Immunofluorescence Staining

Cav1.2, Caveolin-1, PKCα and AKAP150 expression was assessed in isolated cerebral arterial smooth muscle cells, initially fixed in PBS containing 4% paraformaldehyde (15 min, 22 ºC), then washed (3x in PBS), then permeabilized (PBS containing 0.2% Tween 20 (15 min, 22 ºC)) and quenched (PBS containing 0.2% Tween 20 and 3% BSA (22 ºC)). Primary antibodies (**Suppl. Table S1**) diluted in quench solution were then applied overnight (4 ºC); the next day, the smooth muscle cells were washed in PBS (0.2% Tween 20) and treated with fluorophore-conjugated secondary antibodies (Alexa Fluor® 488 donkey anti-rabbit IgG (1:1000) or Alexa Fluor® 488 donkey anti-mouse IgG (1:1000) dependent on primary antibody host species (1 hour, 22 ºC)). Nuclei were stained with Prolong Diamond Antifade Mountant with DAPI, and coverslips were sealed with nail polish. Secondary controls were conducted with the primary antibody removed. Images were captured using a Leica-TCS SP8 or CF-NIKON confocal microscope with a 63× oil-immersion lens and processed using Leica LAS X or MIS-Elements AR analysis software. This staining was used only to define the best antibody dilution for proximity ligation assay (PLA) experiment. Chosen concentrations are listed in **Supplementary Materials Table S1**.

### Proximity Ligation Assay (PLA)

The Duolink *in situ* PLA detection kit assessed the proximity of two proteins in mouse cerebral arterial smooth muscle cells. Enzymatically isolated cells were settled onto cover glass, pretreated with isoosmotic/hypoosmotic solutions and fixed in PBS containing 4% paraformaldehyde (15 min, 22 ºC), washed (3X in PBS) and then exposed to Duolink blocking solution in a heated humidity chamber (1 hour, 37 ºC). Primary antibodies specific to Cav1.2 and caveolin-1, or AKAP150, or PKCα (**Suppl. Table S1**) were mixed with Duolink antibody diluent and applied to cells overnight (4 ºC); controls were completed in parallel with one or both primary antibodies removed. The next day, cells were washed (3X) with Duolink wash buffer and labelled with Duolink PLA plus (rabbit) and minus (mouse) probes (1 hour, 37 ºC). PLA secondary probes are labelled with oligonucleotides that hybridize only when tagged proteins are <40 nm from one another. Following ligation (30 minutes, 37 ºC) and amplification of probe templates (100 minutes, 37 ºC), red fluorophore-tagged complementary oligonucleotides were used to detect protein interactions. Nuclei were stained with Prolong Diamond Antifade Mountant with DAPI and coverslips were sealed with nail polish. Images were captured as described above; ImageJ software was used to count the number of product specific dots per cell. Figures of PLA staining are provided in **Supplementary Materials Fig. S3-6**.

### dSTORM Super-Resolution Microscopy

Isolated smooth muscle cells adherent to a circular coverslip (#1.5) were exposed (10 min) to an isoosmotic (300mOsm, calculated based on chemical composition) (in mM: 82 NaCl, 1 CsCl, 100 nM CaCl_2_, 1.2 MgCl_2_, 10 HEPES, 10 glucose, and 80 mannitol (pH 7.4)) or hypoosmotic (230mOsm, calculated based on chemical composition) (in mM: 82 NaCl, 1 CsCl, 100 nM CaCl_2_, 1.2 MgCl_2_, 10 HEPES, and 10 glucose (pH 7.4)) bath solution. Cells were then fixed and incubated for 20 min at 37° C in glyoxal fixing solution. Following 3 rinses in PBS, coverslips were washed 3x for 5 min with PBS under gentle agitation (30 rpm) and incubated for one hour with blocking solution (0.01% Triton-X in a 1:1 solution of SEA BLOCK:PBS). After removal of the blocking solution, cells were incubated with 0.02 µg/mL custom rabbit anti-FP1 (α1C) 36 in blocking solution overnight at 4° C; custom antibody has been characterized in details previously (39–41). Coverslips were then rinsed and washed (5x for 15 min) with PBS under gentle agitation before being incubated with a secondary antibody (Alexa donkey anti-rabbit 647, 1:1000) in PBS. Cells were rinsed and washed 5x for 15 min in PBS and stored at 4° C until imaging was performed. Secondary antibody controls were performed in experiments where the primary antibody was omitted. Imaging was performed using coverslips mounted on a round cavity microscope slide (Sigma) containing ONI Bcubed proprietary buffers A+B (100:1) (ONI). Images were obtained using the ONI Nanoimager, a super-resolution direct Stochastic Optical Reconstruction Microscopy (dSTORM) microscope coupled to a 100× Olympus oil-immersion UPlanApo TIRF 1.5NA lens. Images were acquired with 10 ms exposure over 30,000 frames with a 640 nm excitation laser and appropriate emission filters. Images were processed and localization maps were obtained using the NimOS localization software (Oxford Nanoimaging). The following filtering parameters were applied to the localization maps during their final rendering: photon count > 600, localization precision (x and y) min: 0 nm, max: 25 nm, sigma (x and y) min: 100 nm, max: 400 nm, p-value min: 0.6, max: 1. All pixels were thresholded, binarized, and segmented into individual objects and included as clusters in our analysis. Cluster size and density were determined using the Analyze Particle option in the ImageJ v2.1.0 (NIH). The BioVoxxel Toolbox plug-in in the ImageJ v2.1.0 (NIH) software was used for neighbor analysis with a neighborhood radius of 250 nm. Frequency distribution histograms of the nearest-neighbor analysis were fitted to Gaussian curves and statistical analysis done with extra sum-of-squares F test comparing the centroids between groups. In initial experiments, our sampling regime was restricted to two regions (2.5 mm^2^) per cell but later expanded to the whole-cell as computational capacity and AI driven algorithmic analysis improved (36). For two color acquisitions, images were acquired simultaneously with 30 ms exposure over 10,000 frames and both 561 and 640 lasers at 50% gain. Localization maps were obtained using the NimOS localization software (Oxford Nanoimaging) and the following filtering parameters were applied for rendering: photon count > 600, localization precision (x and y) min: 0 nm, max: 25 nm, sigma (x and y) min: 100 nm, max: 400 nm, p-value min: 0.6, max: 1. For analysis, all pixels were thresholded, binarized, and segmented into individual objects. Cluster size and density, as well as protein overlap, were determined using ImageJ v2.1.0 (NIH).

### Vessel Myography and Intracellular Ca^2+^ Measurements

Cerebellar and posterior arteries (∼130–180-µm diameter) were stripped of endothelium by passing an air bubble through the lumen; removal was confirmed by the loss of bradykinin-induced dilation. Arterial diameter was monitored by using a ×10 objective and an automated edge detection system (IonOptix); fluorescence changes were captured using RETRA light engine (Lumencor) laser source, ×10 objective, ORCA-Flash4.0 LT plus (Hamamatsu) digital camera, and MIS-Elements AR software.

For pressure myography, de-endothelialized arteries were mounted in a custom arteriograph and superfused with warm (37 ºC) physiological solution (PSS) containing (in mM): 119 NaCl, 4.7 KCl, 20 NaHCO_3_, 1.7 KH_2_PO_4_, 1.2 MgSO_4_, 1.6 CaCl_2_, and 10 glucose. PSS was bubbled with a 5% CO_2_/95% air gas mixture and pH was maintained at ∼7.4. Arteries were equilibrated for 30 minutes at 15 mm Hg and contractile responsiveness assessed by brief exposure (10 s) to 55 mM KCl. Equilibrated arteries were then subjected to 1 of 3 protocols: 1) 20 mmHg pressure steps between 20 to 100 mmHg; 2) administration of 30mM KCl (in PSS) with vessels set at 20 and 80 mmHg to elicit a standardized depolarized; and 3) superfused phenylephrine (10^−9^-to-5*10^−4^ M) with vessels set at 20 mmHg. Vessels’ passive diameter was ascertained in a Ca^2+^-free PSS solution supplemented with 2 mM/L EGTA. Percentage of constriction was calculated using the following formula:

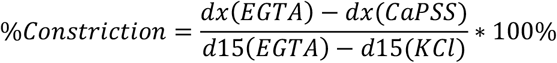

Where *dx(EGTA)* and *dx(CaPSS)* refer to diameter of the vessel (x, µm) at the same pressure in Ca^2+^ free PSS solution (+2 mM/L EGTA) and standard Ca^2+^ containing PSS, respectively. *d*15 refers to diameter of the vessel at 15 mmHg. When necessary, wall stress (σ, kPa) was calculated and considered proportional to the transmural pressure (P, kPa) and radius (R, m) and inversely proportional to the average vessel wall thickness (w, m):

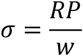

To assess cytosolic [Ca^2+^], isolated cerebral arteries were incubated in PBS containing 5 μM Fura-2AM (1 h, 4ºC). Fura-2AM is a ratiometric probe designed to detect changes in [Ca^2+^]_i_ as a ratio between excitation/emission of 340/510 nm (measure of Ca^2+^-bound Fura-2AM) and 380/510 nm (measure of Ca^2+^-free Fura-2AM). Dye-loaded arteries were then mounted in a custom arteriograph, superfused with warm (37 ºC) PSS (5% CO_2_/95% air, pH=7.4) and contractile responsiveness assessed by brief exposure (10 s) to 55 mM KCl (15 mmHg). Following equilibration, arteries were then pressurized to 20 or 80 mmHg and the changes in [Ca^2+^]_i_ measured at rest and with the vessels depolarized to same extent via administration of 30mM KCl **(Suppl. Figure S7)**. At 20 mmHg with 30 mM KCl, the change in [Ca^2+^]_i_ was exclusively due to V_m_-dependent opening of Ca_V_1.2 channels; while at 80 mmHg with 30mM KCl, the change in [Ca^2+^]_i_ was higher and represented an integrated response of both V_m_-dependent opening of Ca_V_1.2 channels and functional coupling/perimembrane trafficking triggered by pressure stimuli.

### Computational Analysis

An in-house developed algorithm was used to process the cell-attached Ca_V_1.2 recordings and to calculate key coupling parameters. Each recording was first detrended to remove any potential slow drift and then filtered using an 8-point Bessel filter with a suitable cutoff frequency. Single Ca_V_1.2 opening current was assumed as q = 0.5 pA based on literature and the content of the patch pipette solution (42). Following published approaches (43), the values of the detrended and filtered recording at each time point were then binned using integer multiples of q, which provided a simple yet reproducible idealized trace. The maximum number of channels, N, was assumed to be the largest integer multiple of q that was found in the idealized trace. The idealized trace was used to estimate a data driven matrix of transition probabilities. Since the channels in the patch-clamp are identical, a theoretical transition probability matrix of the aggregated cluster was constructed as described in literature (44). Using the two matrices, the single independent channel open to open transition probability (r), single independent channel closed to closed transition probability (z), and channel-channel coupling parameter (k) were estimated. The estimation was performed using a standard Nelder-Mead simplex procedure, see (45) for details. In addition to identifying the underlying Markov chain, the idealized trace was analyzed for other channel statistics which consisted of the probability of one or more channels being open (nP_o_). The parameter κ, together with N and nP_o_ was used as quantitative measures of the potential effects of pressure on Cav1.2 behavior.

### In silico model describing microvascular flow and cerebral autoregulation

A brief overview of the *in silico* model designed by Lambride et al. is provided herein (17). The workflow typically advances in three main steps: (1) generation of microvascular networks (MVNs), (2) simulation of cerebral blood flow (CBF) with autoregulatory control, and (3) analysis of simulation outputs. The MVN is derived from 3D maximal projections (two-photon fluorescent microscopy) of surface arterial (SA) vessels of a BALB/c mouse (**Fig. 9A**). The network covers a 3.5 × 3.5 mm^2^ region of the cortical surface in the whisker and hindlimb areas, capturing branches from the anterior and middle cerebral artery (ACA, MCA) territories (MCA-M4/M5). After manual tracing and conversion into graph format, descending arteriole (DA) root density was increased to match *in vivo* observations. To complete the cortical microvascular architecture, DA and ascending venule (AV) trees were added and connected via an artificial capillary bed (Cs). Additional arteries were included to connect MCA- and ACA-sided SAs to a common inflow node, coming from the circle of Willis (CoW). To reduce uncertainties and align the flow field to *in vivo* data, we calibrated vessel diameters using an inverse modeling approach(46, 47). The resulting MVN serves as the baseline configuration. A detailed description of the generation and validation of this approach is provided in Epp et al. (47).

The *in silico* blood flow model was derived from Poiseuille’s law and mass conservation at bifurcations (47, 48). We assumed a constant tube hematocrit of 0.3 and accounted for the Fahraeus-Lindqvist effect *via* an empirical formulation (*in vitro* version) (49). The MVN is characterized by a single inflow vertex at the most upstream SAs connected to CoW, and multiple outflow vertices at the roots of AVs. Pressure boundary conditions were applied as 100 mmHg at the CoW (representing mean arterial pressure, MAP) and 10 mmHg at the AV outlets (47).

Blood vessels were modeled as elastic tubes whose diameters adapt in response to changes in pressure through passive and active mechanisms. Passive responses are governed by the intrinsic mechanical properties of the vessel wall, resulting in dilation when pressure rises and constriction when pressure falls. Active regulation reflects physiological control mechanisms that adjust vascular smooth muscle (VSM) tone to maintain CBF stability in response to changes in MAP (50, 51). In our model we consider two key mechanisms, namely myogenic and endothelial responses. For both passive and active mechanisms, we used a pressure–area relationship based on linear elastic theory, where transmural pressure governs changes in vessel cross-sectional area (52). Passive (non-regulating) vessels are AVs and Cs, that respond solely to changes in transmural pressure. Actively regulating vessels, including all arteries (SAs and DAs). These vessels adapt their compliance in response to physiological stimuli, thereby modifying the pressure–area relationship (53, 54).

To model active regulatory adaptation, we used a quasi-steady feedback formulation that incorporates both myogenic and endothelial responses. Following Daher et al. (54), the relative vessel wall stiffness *k*_*ij*_ is given by:

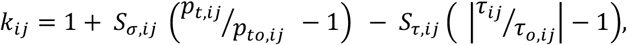

where *S*_*σ,ij*_ and *S*_*τ,ij*_ are sensitivity factors for myogenic and endothelial responses, respectively. The myogenic component is driven by changes in transmural pressure *p*_*t,ij*_, while, the endothelial response depends on changes in wall shear stress *τ*_*ij*_.The baseline values, *p*_*to,ij*_ and *τ*_*o,ij*_, were computed using baseline flow and pressure fields, i.e., the configuration directly obtained after employing the inverse model. Based on prior experimental findings, the myogenic mechanism was modeled as the dominant component, with *S*_*σ,ij*_ typically larger than *S*_*τ,ij*_, which is reflected by the sensitivity factors of 4.0 and 0.5, respectively. As compliance can only vary within set limits from its baseline value, we applied a sigmoidal function to link the stiffness *k*_*ij*_ to relative compliance *C*_*r,ij*_ (54):

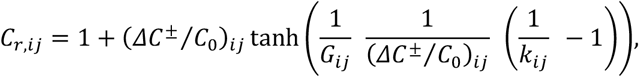

where (Δ*C*^±^/*C*_0_)_*ij*_ defines the maximum bounds of positive and negative compliance changes (asymmetrically constrained) (54, 55) and *G*_*ij*_ sets the steepness of the compliance-stiffness relationship.

Eventually, changes in compliance modify the pressure–area relation, allowing the vasculature to adapt to pressure variations and reach a new equilibrium. To simulate this adaptative behavior under external stimuli (e.g., MAP changes), we used an iterative approach (**Fig. 9B**). At each iteration, the blood flow model was solved to compute the updated pressure and flow fields while enforcing mass conservation at each node. For regulating vessels, we then computed transmural pressure and shear stress to update stiffness and compliance and used these values to determine the new diameters. The system was solved iteratively until convergence, defined as a maximum relative diameter change below 10^−6^.

In this study, we focused on arterial constriction in response to increased pressure. Specifically, we varied MAP within the range of 100–140 mmHg, with 100 mmHg set as the baseline. The model interprets these variations as relative changes from baseline and decreases vessel diameters through autoregulatory mechanisms to maintain constant flow. To examine how a partial loss of the myogenic response affects arterial constriction and capillary perfusion, we reduced the initial myogenic sensitivity factor (*S*_*σ*_=4.0) by 30% and 50% (*i*.*e*., *S*_*σ*_= 2.8 and 2.0, respectively). The endothelial sensitivity factor was kept constant for all simulations.

We conducted a detailed vessel-level analysis to evaluate how arteries responded to MAP elevations under each sensitivity condition by computing the mean relative change in diameter compared to baseline (MAP = 100 mmHg). As we have previously demonstrated (17) that arterial responses depend on their location, we grouped vessels based on their baseline intraluminal pressure, which serves as a proxy to reflect their position within the network. To assess the perfusion at the capillary level in response to elevated pressure, we computed the relative change in flow rate for each capillary segment *ij* compared to baseline, defined as: 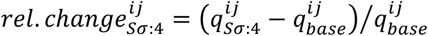. This metric was also computed for the reduced sensitivity factors (*i*.*e*., *S*_*σ*_= 2.8 and 2). To quantify the effect of impaired myogenic response on capillary flow, we calculated the median relative difference between impaired and healthy conditions using *median*(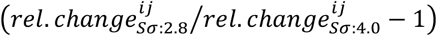). Using the same metrics, we examined the flow changes in capillaries across cortical depths. Additional information can be found in **Supplemental materials Fig. S8**.

### Statistical Analysis

Data were analyzed using GraphPad Prism v10 and presented as means ± standard error of the mean. P-values < 0.05 were considered statistically significant, the exact p-values are reflected on the figures. The two-way ANOVA statistical test was used for whole-cell patch-clamp experiments, as it included two changing parameters, i.e., voltage and osmolarity. The rest of the data was analyzed using unpaired t-test when comparing summary data of cohorts of control and experimental data (e.g., IHC results), while for matching data sets paired t-test was applied. More detailed information concerning statistical analysis is provided in the figure legend and **Supplementary Table S2**. An a priori power analysis was performed to determine that the planned sample size would yield a statistical power >0.8 at α = 0.05, based on the expected effect size from pilot data.

## ABBREVIATIONS

PKC: protein kinase C
PKA: protein kinase A
AKAP: A kinase associated protein (called AKAP150 in rodents and AKAP79 in human)
Noc: Nocodazole
Calph C: Calphostin C
Cav1/Cav1 SDP: caveolin-1 scaffolding domain peptide and its scrambled control
ATP: adenosine triphosphate
HEPES: 4-(2-hydroxyethyl)-1-piperazineethanesulfonic acid
EGTA: ethylene glycol-bis(β-aminoethyl ether)-N,N,N′,N′-tetraacetic acid
PBS: phosphate-buffered saline
PSS: physiological saline solutions
TIRF: total internal reflection fluorescence microscopy
DAPI: 4′,6-diamidino-2-phenylindole
SMC: smooth muscle cell
[Ca^2+^]_i_: intracellular concentration of Ca^2+^
Vm: membrane potential

## Sources of Funding

Supported by the Canadian Institutes for Health Research and the Rorabeck Chair in Vascular Biology and Neuroscience, CL & FS received funding from the Swiss National Science Foundation (Grant No. 200703 and 202199) and the Hartmann Müller Foundation (Grant No. 2885).

## Acknowledgments

The authors acknowledge the valuable help of Muhammad Zahoor, and excellent technical support of Suzanne E. Brett and Michelle S. Kim (University of Western Ontario). The schematic diagrams on the figures (except Fig. 9) were created using BioRender.com

## Data availability

The data that support the findings of this study are available from the corresponding author upon reasonable request.

## Code availability

The in-house algorithm (Markov chains) can be found at https://github.com/mccsssk2/Mironova_et_al_20242025\

A detailed description of the *in silico* model is available in Lambride, C., et al. (2025) (17), Loss of Cerebral Autoregulation After Stroke Drives Abnormal Perfusion Patterns.The simulation code is available at https://github.com/Franculino/microBlooM (v2.0.0).

## Disclosures

None.

## Notes

https://github.com/mccsssk2/Mironova_et_al_20242025\

https://github.com/Franculino/microBlooM

### Competing Interest Statement

The authors have declared no competing interest.

