## Supplemental tables 1-2, Supplemental figures 1-11 for "Blood Pressure Regulates Functional Coupling of L-Type Ca^2+^ Channels: Reimaging the Foundation of Cerebral Blood Flow Control"

**Correspondence:**

Supporting text
Figures S1 to S11
Tables S1 to S2

**Section A. Supplementary table.**

**Table S1. Primary and Secondary Antibody Characteristics**

| Antibody (Ab) | Raised against amino acids (aa)/ antigen (Ag, species / epitope) | [Ab] | Supplier/Cat # | Host | Application |
|---|---|---|---|---|---|
| Anti-CACNA1C ($Ca_V1.2$) | Recombinant Protein within Rabbit CACNA1C aa 1500-1750 | 1:50 | Abcam/ab84814 | Mouse | IHC, PLA |
| Anti-Cav-1 | Synthetic peptide corresponding to a region at the N-terminus of human caveolin-1 (amino acids 2-20) | 1:200 | Sigma/C4490 | Rabbit | IHC, PLA |
| Anti-AKAP150 | Peptide corresponding to amino acids 428-449 of rat AKAP 150 (C-TTVGQAEEATVGQAEEATVGQA) | 1:200 | EMD Millipore/07-210 | Rabbit | IHC, PLA |
| PKCα | Synthesized peptide derived from human PKC alpha. | 1:200 | Sigma/SAB4502354 | Rabbit | IHC, PLA |
| Alexa Fluor® 488 donkey IgG | Rabbit IgG(H+L) | 1:1000 | ThermoFisher/A21206 | Donkey | IHC |
| Alexa Fluor® 488 donkey IgG | Mouse IgG(H+L) | 1:1000 | ThermoFisher/A21202 | Donkey | IHC |
| Alexa Fluor® 647 donkey IgG | Rabbit IgG(H+L) | 1:1000 | ThermoFisher/A31573 | Donkey | SR |
| Custom rabbit anti-FP1 (α1C) | Rabbit | 0.02 µg/mL | NA | NA | SR |

Application key: IHC - immunohistochemistry, PLA - proximity ligation assay, SR – super resolution microscopy

**Table S2. Statistical information for the manuscript figures**

| Figure # | Statistical method | N of animals/cells | Additional information |
|---|---|---|---|
| Figure 1B | 2way ANOVA, Sidak's multiple comparison test | 6 cells from 6 animals | There was a significant main effect of voltage, $F(9, 50) = 25$, $p < 0.0001$; pressure, $F(1, 50) = 31$, $p < 0.0001$; voltage × pressure |

| | | | |
|---|---|---|---|
| | | | interaction, F(9, 50) = 2.5, p = 0.0189. |
| Figure 1D | 2way ANOVA, Sidak's multiple comparison test | 7 cells from 7 animals | There was a significant main effect of voltage, $F(9, 60) = 7.3$, $p < 0.0001$; no significant main effect of pressure, $F(1, 60) = 0.0$, $p = 0.8126$; the voltage × pressure interaction was not significant, $F(9, 60) = 0.1$, $p = 0.9978$. |
| Figure 1E left | 2way ANOVA, Sidak's multiple comparison test | 5 cells from 5 animals | There was a significant main effect of voltage, $F(9, 40) = 10$, $p < 0.0001$; pressure, $F(1, 40) = 50$, $p < 0.0001$;t voltage × pressure interaction, $F(9, 40) = 3.8$, $p = 0.0015$. |
| Figure 1E right | 2way ANOVA, Sidak's multiple comparison test | 5 cells from 5 animals | There was a significant main effect of voltage, $F(9, 40) = 17$, $p < 0.0001$; pressure, $F(1, 40) = 36$, $p < 0.0001$; voltage × pressure interaction, $F(9, 40) = 3.4$, $p = 0.0032$. |
| Figure 1F | Unpaired t test with Welch's correction, one-tailed hypothesis | N of animals equals n of cells, left to the right: 6, 5, 7, 6, 5, 7 | Left to the right: t=2.177, df=8.924; t=2.263, df=7.607; t=0.9028, df=6.086; t=1.862, df=11.00. |
| Figure 2B | Wilcoxon matched-pairs signed rank test, one-tailed hypothesis | 6 cells from 6 animals | W= -21.00 |
| Figure 2C | Wilcoxon matched-pairs signed rank test, one-tailed hypothesis | 6 cells from 6 animals | W= -21.00 |
| Figure 2E | Wilcoxon matched-pairs signed rank test, one-tailed hypothesis | 6 cells from 6 animals | W= -5.00 |

| | | | |
|---|---|---|---|
| Figure 2F | Wilcoxon matched-pairs signed rank test, one-tailed hypothesis | 6 cells from 6 animals | W= -3.00 |
| Figure 2H | Wilcoxon matched-pairs signed rank test, one-tailed hypothesis | 6 cells from 6 animals | W= 0.00 |
| Figure 2I | Wilcoxon matched-pairs signed rank test, one-tailed hypothesis | 6 cells from 6 animals | W= -14.00 |
| Figure 3B left | 2way ANOVA, Sidak's multiple comparison test | 6 cells from 6 animals | There was a significant main effect of voltage, $F(9, 50) = 25$, $p < 0.0001$; pressure, $F(1, 50) = 31$, $p < 0.0001$; voltage × pressure interaction, $F(9, 50) = 2.5$, $p = 0.0189$. |
| Figure 3B right | 2way ANOVA, Sidak's multiple comparison test | 7 cells from 7 animals | There was a significant main effect of voltage, $F(9, 50) = 18$, $p < 0.0001$; pressure, $F(1, 50) = 5.4$, $p = 0.0239$; the voltage × pressure interaction was not significant, $F(9, 50) = 1.6$, $p = 0.1167$. |
| Figure 3C | 2way ANOVA, Sidak's multiple comparison test | 6 cells from 6 animals | There was a significant main effect of voltage, $F(9, 50) = 9.0$, $p < 0.0001$; pressure, $F(1, 50) = 5.5$, $p = 0.0228$; voltage × pressure interaction, $F(9, 50) = 6.7$, $p < 0.0001$. |
| Figure 3D | 2way ANOVA, Sidak's multiple comparison test | 6 cells from 6 animals | There was a significant main effect of voltage, $F(9, 45) = 11.1$, $p < 0.0001$; there was no significant effect of pressure, $F(1, 5) = 1.3$, $p = 0.3055$; and no significant voltage × pressure interaction, $F(9, 45) = 0.6$, $p = 0.7486$. |

| | | | |
|---|---|---|---|
| Figure 3G | Wilcoxon matched-pairs signed rank test, one-tailed hypothesis | 9 cells from 9 animals | W=13.00 |
| Figure 3H | Wilcoxon matched-pairs signed rank test, one-tailed hypothesis | 9 cells from 9 animals | W=15.00 |
| Figure 4B | Mann-Whitney test, two-tailed hypothesis | 50 cells for isosmotic solution, 59 cells for hypoosmotic solution from 3 animal | Overlapping: U= 617<br>Non-overlapping: U=895 |
| Figure 4C | Mann-Whitney test, two-tailed hypothesis | 49 cells for isosmotic solution, 59 cells for hypoosmotic solution from 3 animal | U=758 |
| Figure 4D | Mann-Whitney test, two-tailed hypothesis | 16 cells for isosmotic solution, 20 cells for hypoosmotic solution from 3 animal | U=89 |
| Figure 4E | Mann-Whitney test, two-tailed hypothesis | 20 cells for isosmotic solution, 20 cells for hypoosmotic solution from 3 animal | U=155 |
| Figure 4F | Mann-Whitney test, two-tailed hypothesis | 14 cells for isosmotic solution, 17 cells for hypoosmotic solution from 3 animal | U=68 |
| Figure 4G | Mann-Whitney test, two-tailed hypothesis | 16 cells for isosmotic solution, 20 cells for hypoosmotic solution from 3 animal | U=113 |
| Figure 4H | Mann-Whitney test, two-tailed hypothesis | 20 cells for isosmotic solution, 21 cells for hypoosmotic solution from 3 animal | U=72 |
| Figure 4I | Mann-Whitney test, two-tailed hypothesis | 14 cells for isosmotic solution, 18 cells for hypoosmotic solution from 3 animal | U=25 |
| Figure 5B Control | Mann-Whitney test, two-tailed hypothesis | 27 cells for isosmotic solution, 27 cells for hypoosmotic solution from 3 animal | U=164 |

| | | | |
|---|---|---|---|
| Figure 5B Nocodazole | Mann-Whitney test, two-tailed hypothesis | 19 cells for isosmotic solution, 22 cells for hypoosmotic solution from 3 animal | U=204 |
| Figure 5B Calphostin C | Mann-Whitney test, two-tailed hypothesis | 14 cells for isosmotic solution, 21 cells for hypoosmotic solution from 3 animal | U=98 |
| Figure 5C Control | Mann-Whitney test, two-tailed hypothesis | 27 cells for isosmotic solution, 28 cells for hypoosmotic solution from 3 animal | U=251 |
| Figure 5C Nocodazole | Mann-Whitney test, two-tailed hypothesis | 19 cells for isosmotic solution, 22 cells for hypoosmotic solution from 3 animal | U=136 |
| Figure 5C Calphostin C | Mann-Whitney test, two-tailed hypothesis | 15 cells for isosmotic solution, 21 cells for hypoosmotic solution from 3 animal | U=145 |
| Figure 6H | Wilcoxon matched-pairs signed rank test, one-tailed hypothesis | 16 cells from 16 animals | W=136.00 |
| Figure 6I left to right | Wilcoxon matched-pairs signed rank test, one-tailed hypothesis | 5 cells from 5 animals; 6 cells from 6 animals; 5 cells from 5 animals | W=-15.00; W=-21.00; W=-15.00 |
| Figure 7E right pair | Mann-Whitney test, one-tailed hypothesis | 6 cells from 6 animals | U=0 |
| Figure 7F | 2way ANOVA, Sidak's multiple comparison test | 6 cells from 6 animals | There was a significant main effect of pressure, $F(4, 50) = 8.9$, $p < 0.0001$; myogenic tone, $F(1, 50) = 8.3$, $p = 0.0058$; the pressure × myogenic tone interaction was not significant, $F(4, 50) = 0.1$, $p = 0.9476$. |
| Figure 7H | Wilcoxon matched-pairs signed rank test, one-tailed hypothesis | 7 cells from 7 animals | W=28.00 |

| | | | |
|---|---|---|---|
| Figure 7I | Wilcoxon matched-pairs signed rank test, one-tailed hypothesis | 7 cells from 7 animals | W=28.00 |
| Figure 7K | Wilcoxon matched-pairs signed rank test, one-tailed hypothesis | 7 cells from 7 animals | W=2.00 |
| Figure 7L | Wilcoxon matched-pairs signed rank test, one-tailed hypothesis | 7 cells from 7 animals | W=14.00 |
| Figure 8C | Wilcoxon matched-pairs signed rank test, one-tailed hypothesis | 5 cells from 5 humans | W=15.00 |
| Figure 8E | Wilcoxon matched-pairs signed rank test, one-tailed hypothesis | 8 cells from 8 humans | W=28.00 |
| Figure 8F | Wilcoxon matched-pairs signed rank test, one-tailed hypothesis | 8 cells from 8 humans | W=36.00 |

**Section B. Supplemental figures.**

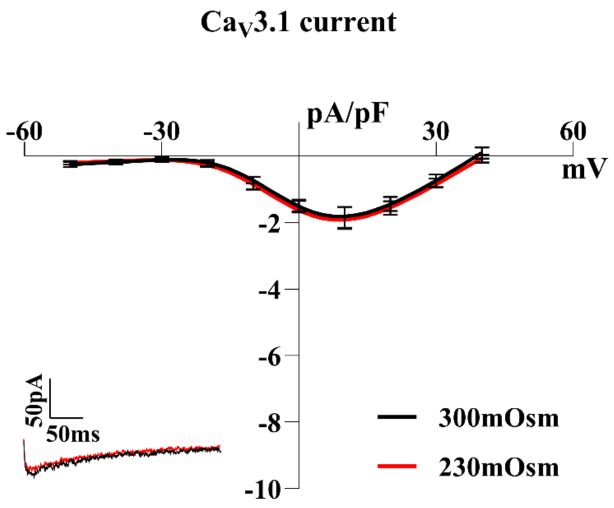


**Figure S1.** Whole-cell Ca$_V$3.1 current was unaffected by the hypoosmotic challenge; Ca$_V$1.2 and Ca$_V$3.2 channels were blocked by 200nM Nifedipine and 50 μM of $Ni^{2+}$, respectively. n=7 cells from 7 animals, two-way ANOVA, Sidak's multiple comparison test.

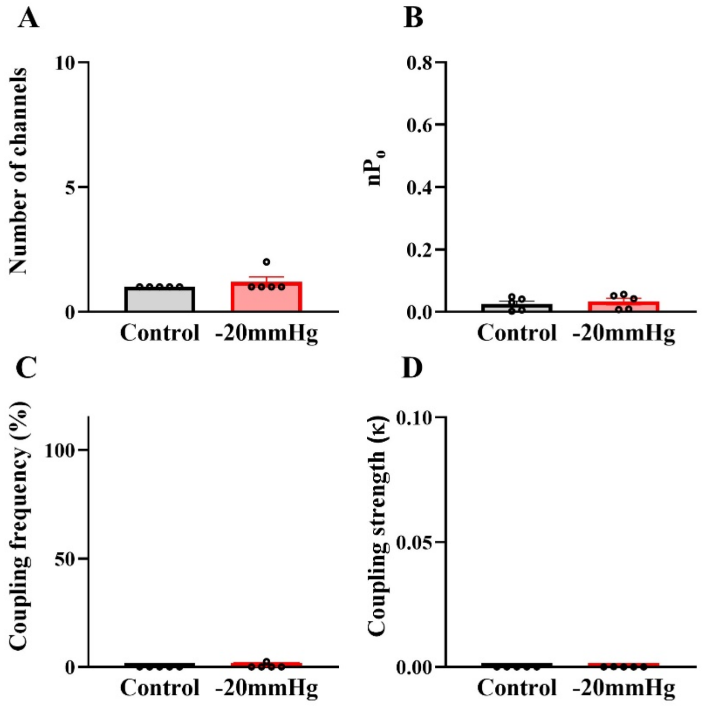


**Figure S2.** Summative single-channel data demonstrates that the $Ca^{2+}$ blocker Nifedipine (200nM in pipette and bath) inhibits L-type $Ca^{2+}$ channels (cell-attached patch-clamp). Number of channels opening (**A**) and nP$_o$ (**B**) were significantly reduced; no functional coupling was observed is these particular experiments (**C&D**). 5 cells from 5 animals, Wilcoxon matched-pairs signed-rank test with a one-tailed hypothesis.

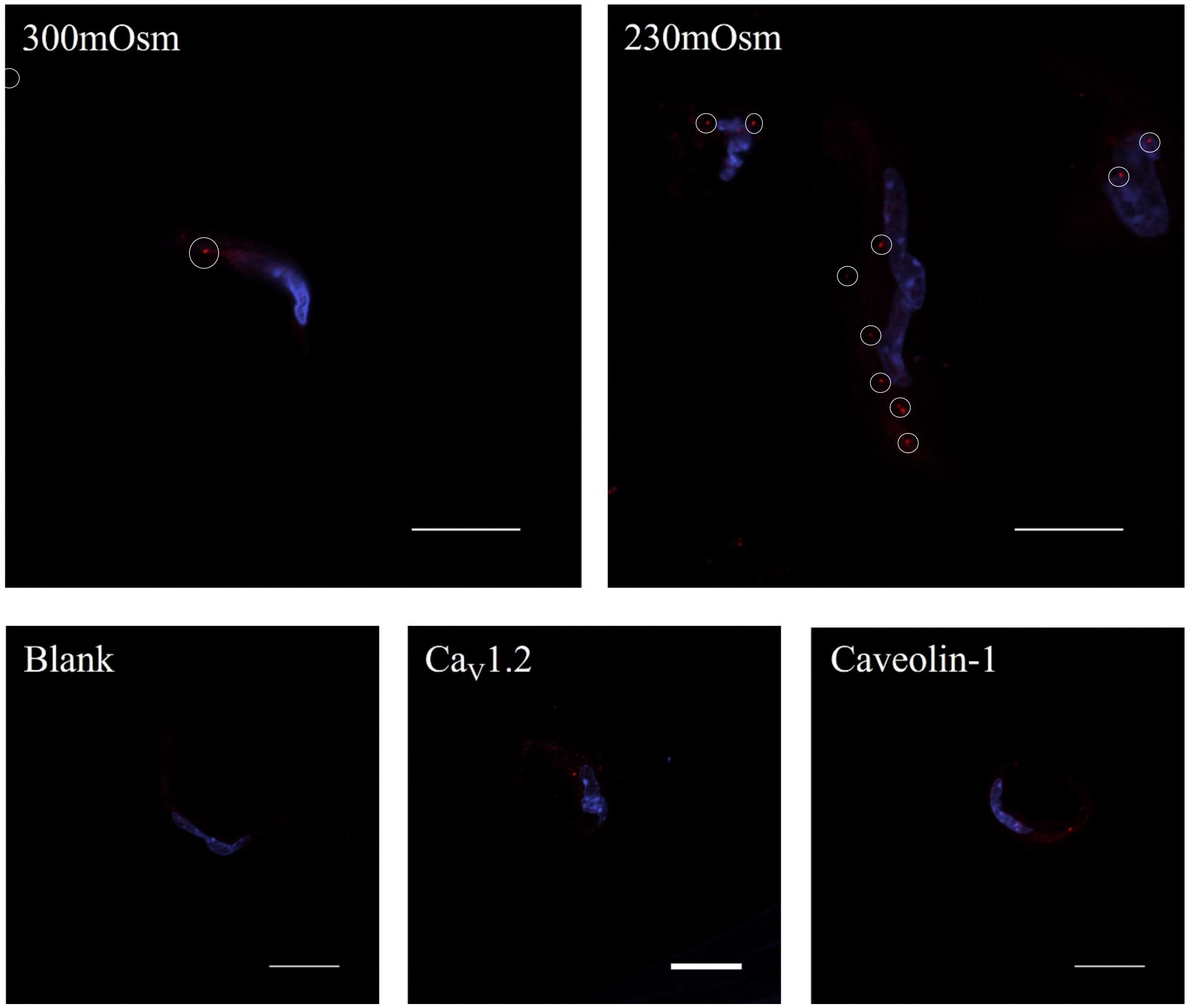


**Figure S3.** Proximity ligation assay reveals enhanced colocalization of $Ca_V1.2$ with Caveolin-1 when cerebral arterial smooth muscle cells are exposed to a hypoosmotic challenge; representative figure with controls. White bar equals 20 micrometers.

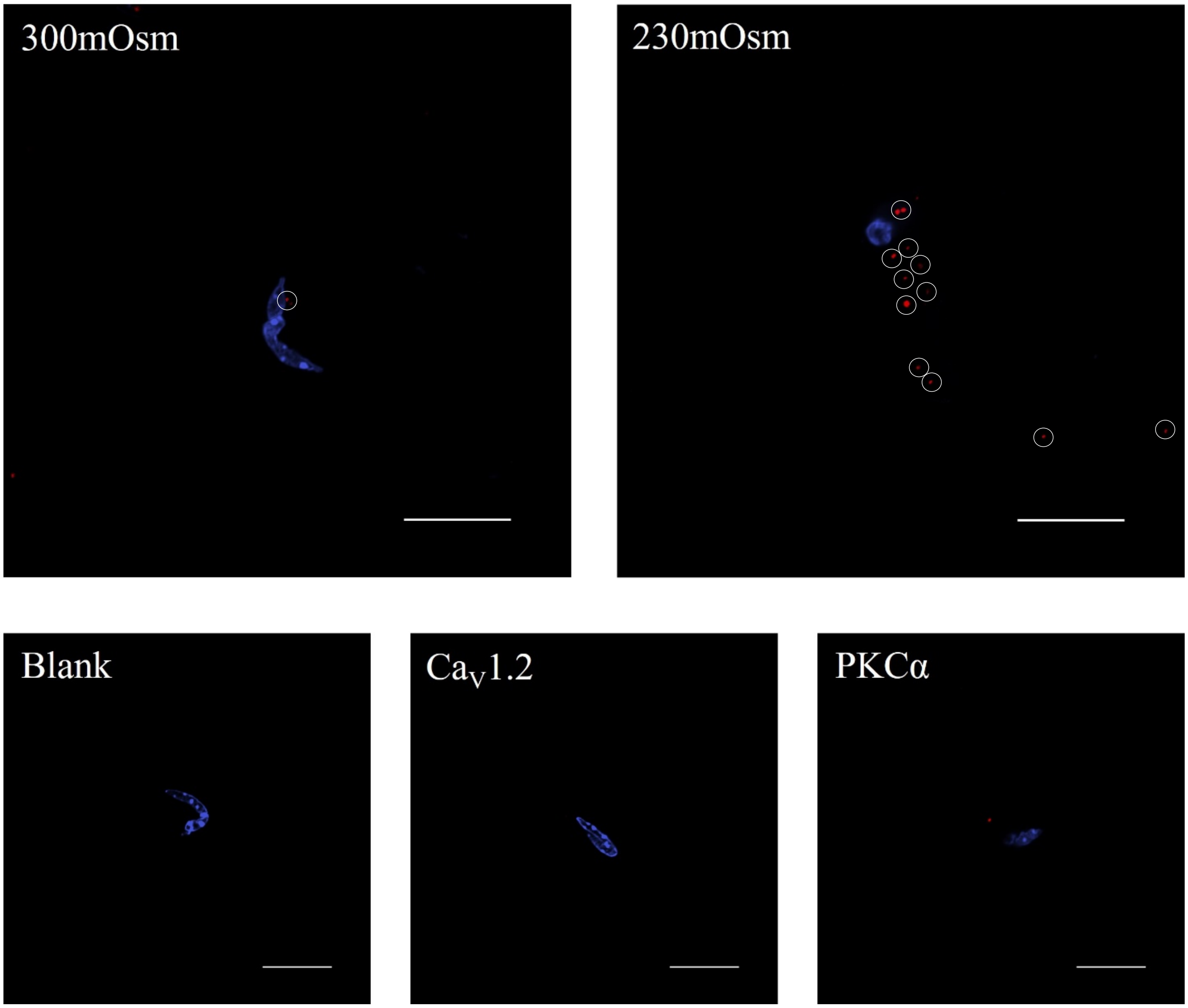


**Figure S4.** Proximity ligation assay reveals enhanced colocalization of $Ca_V1.2$ with PKCα when cerebral arterial smooth muscle cells are exposed to a hypoosmotic challenge; representative figure with controls. White bar equals 20 micrometers.

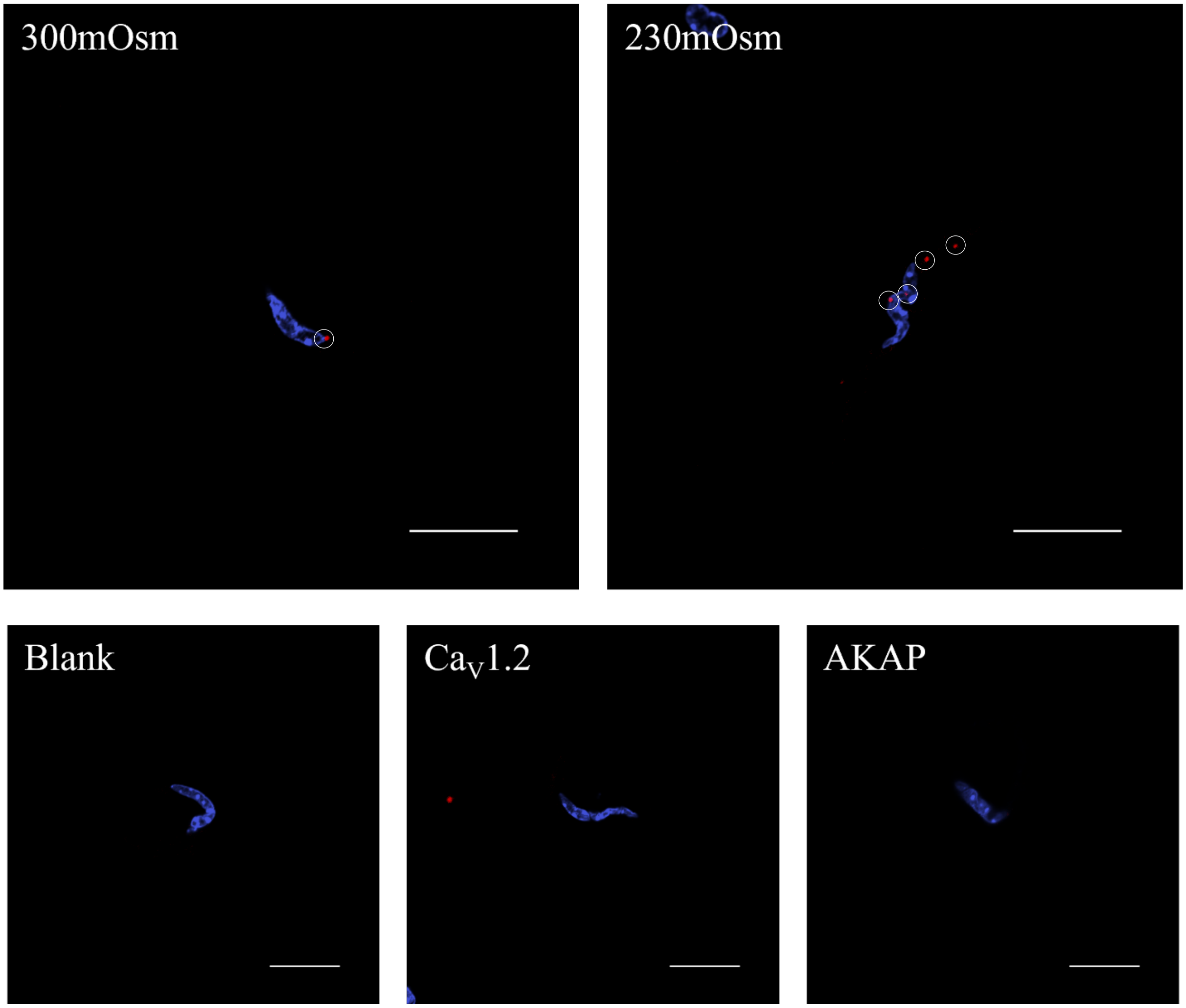


**Figure S5.** Proximity ligation assay reveals enhanced colocalization of $Ca_V1.2$ with AKAP when cerebral arterial smooth muscle cells are exposed to a hypoosmotic challenge; representative figure with controls. White bar equals 20 micrometers.

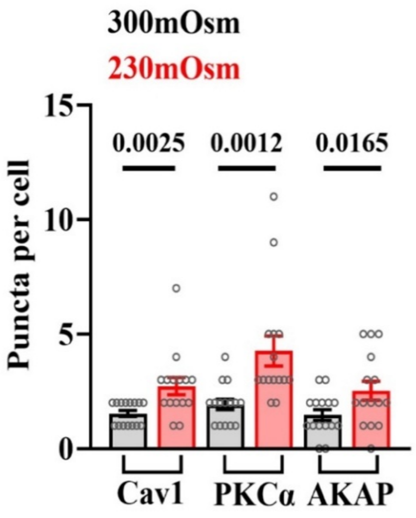


**Figure S6.** Proximity ligation assay reveals that when cerebral arterial smooth muscle cells are exposed to a hypoosmotic challenge, colocalization of $Ca_V1.2$ with AKAP, PKCα, and Caveolin-1 increases (unpaired t-test with one-tailed hypothesis, n=15 cells from 3 animals (5 cells per animal), df=28, left-to right: t=3.043, 3.35, 2.244).

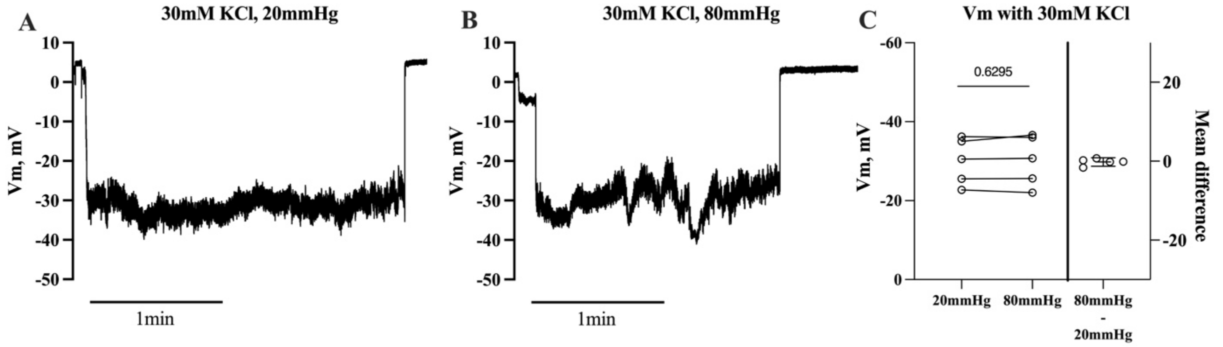


**Figure S7.** Representative traces (**A & B**) and summary data (**C**) show that in the presence of 30 mM KCl, a strong and consistent depolarization was observed in endothelium-denuded cerebral arteries; this depolarization averaged ~-30 mV irrespective of the intravascular pressure. Data was analyzed by paired t test (two tailed); one artery was used per animal animals (n=5 arteries from 5 animals, t=0.5216, df=4). Smooth muscle membrane potential (Vm) was assessed by inserting a glass microelectrode backfilled with 1 M KCl (tip resistance = 100-150 MΩ) into the vessel wall. Measurements of Vm were first made at 20 mmHg in presence of 30mM KCl, and then at 80 mmHg in presence of 30mM KCl in the same artery. The criteria for successful cell impalement included: 1) a sharp negative Vm deflection upon insertion; 2) a stable recording for at least 1 min following entry; and 3) a sharp return to baseline upon electrode removal.

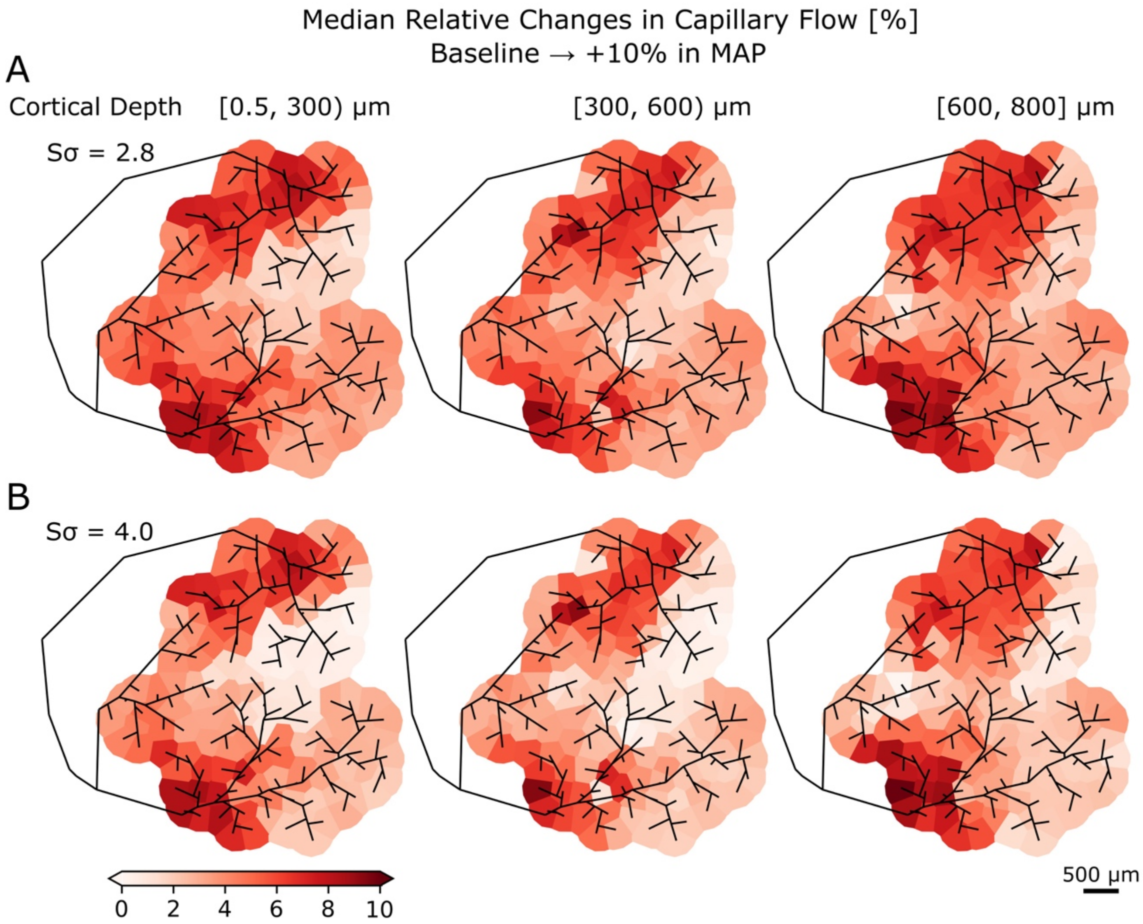


**Figure S8.** Capillary perfusion change in response to a 10% increase in mean arterial pressure (MAP) at two different "myogenic sensitivities". Capillaries are grouped in columns below each descending artery (DA) root point to allow a planar representation. Each panel shows the median relative change in capillary flow between baseline MAP and a 10% increase, for Sσ = 2.8 **(A)** and Sσ = 4.0 **(B)**, at three cortical depth ranges: [0.5, 300), [300, 600], and [600, 800] μm.

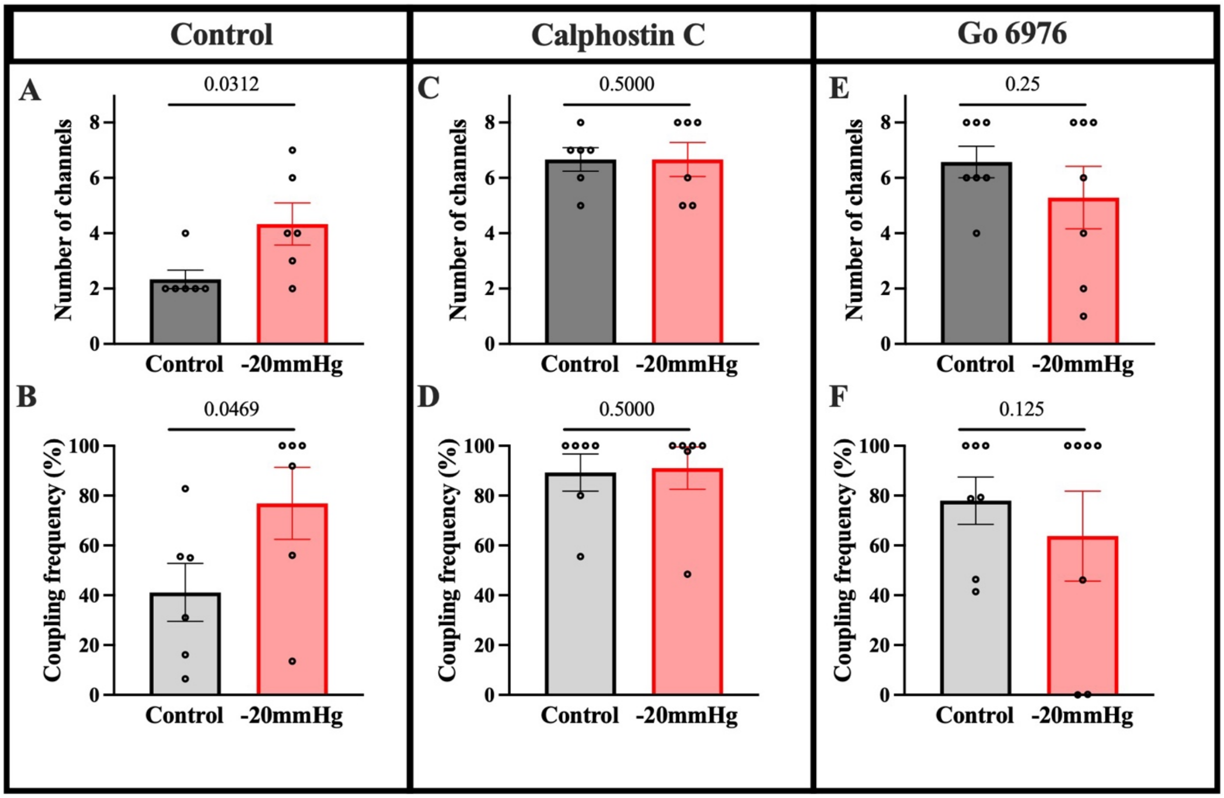


**Figure S9.** Mechano-stimulation increases $Ca_V1.2$ single-channel activity (mouse cerebral arterial smooth muscle cells) in a PKC dependent manner. Statistical analysis of single-channel recordings of $Ca_V1.2$ activity prior to and following negative pipet pressure application (-20 mmHg) under control conditions (**A-B**), in presence of Calphostin C (300nM; broad spectrum PKC inhibitor (**C-D**)) and Go 6976 (100nM; PKCα selective inhibitor (**E-F**)). Analysis revealed that negative pressure application increased number of $Ca_V1.2$ channels and their coupling frequency (**A-B**), (6 cells from 6 animals, Wilcoxon matched-pairs signed-rank test with a one-tailed hypothesis). Calphostin C or Go 6976 both inhibited increase of number of channels (**C, E**) and coupling frequency (**D, F**), (6 cells from 6 animals for Calphostin C and 7 cell from 7 animals for Go 6976; Wilcoxon matched-pairs signed-rank test with a one-tailed hypothesis).

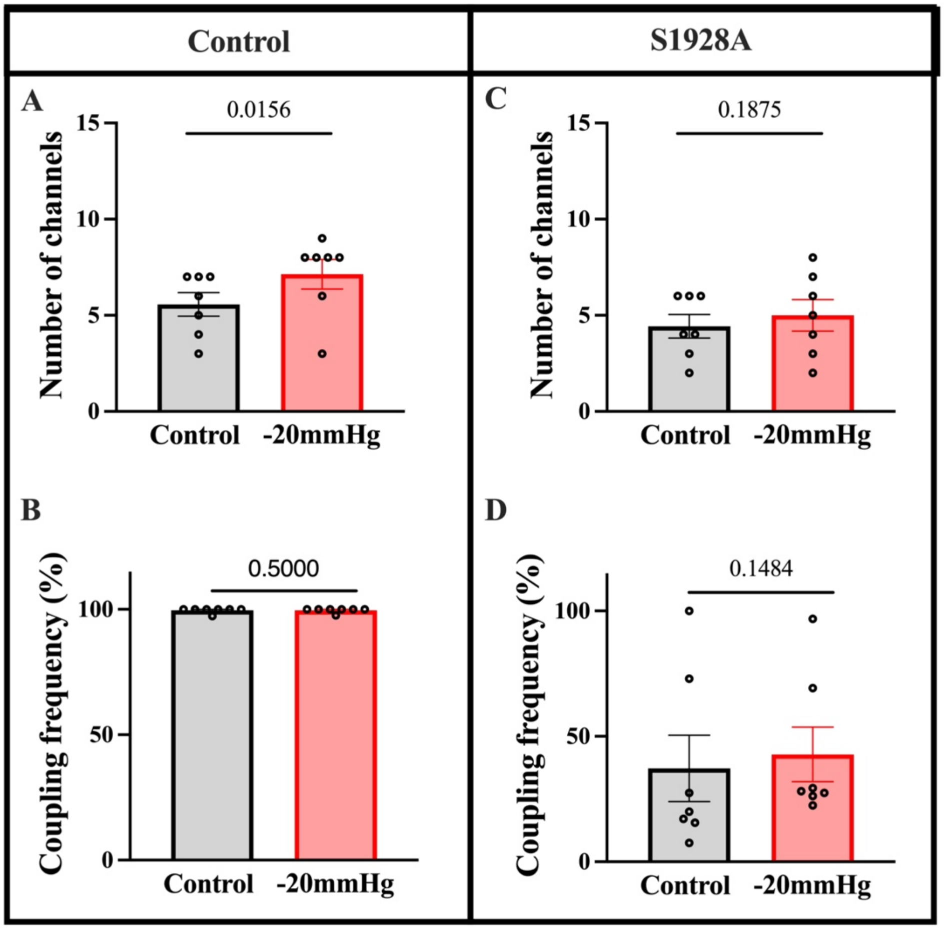


**Figure S10.** Statistical analysis of single-channel $Ca_V1.2$ activity prior to and following negative pipet pressure application (-20 mmHg) in control C57 (**A-B**), and in S1928A (**C-D**) mice revealed that cerebral arterial smooth muscle cells from S1928A mice were insensitive to mechano-stimulation, specifically the number of channels or coupling frequency did not significantly change (n in both control and S1928A mice: 7 cell from 7 animals; Wilcoxon matched-pairs signed-rank test with a one-tailed hypothesis).

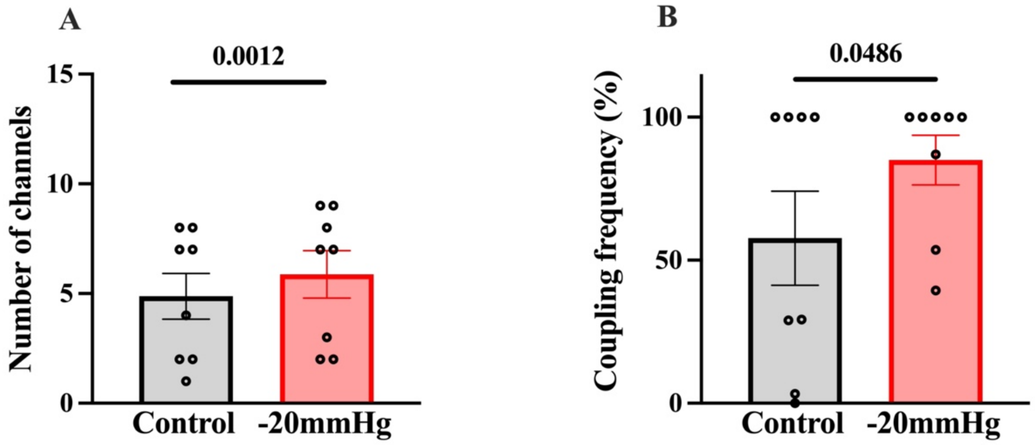


**Figure S11.** Statistical analysis of single-channel recordings of $Ca_V1.2$ activity prior to and following negative pipet pressure application (-20 mmHg) under control conditions demonstrates that negative pressure application increased number of $Ca_V1.2$ channels (**A**) and coupling frequency (**B**). 8 cells from 8 humans; Wilcoxon matched-pairs signed-rank test with a one-tailed hypothesis.
